# Cryo-EM structure of C9ORF72-SMCR8-WDR41 reveals the role as a GAP for Rab8a and Rab11a

**DOI:** 10.1101/2020.04.16.045708

**Authors:** Dan Tang, Jingwen Sheng, Liangting Xu, Xiechao Zhan, Jiaming Liu, Jiang Hui, Xiaoling Shu, Xiaoyu Liu, Tizhong Zhang, Lan Jiang, Cuiyan Zhou, Wenqi Li, Wei Cheng, Zhonghan Li, Kunjie Wang, Kefeng Lu, Chuangye Yan, Shiqian Qi

**Author notes:** These authors contributed equally to this work (Dan Tang, Jingwen Sheng, Liangting Xu). To whom correspondence may be addressed. or (ORCID: 0000-0002-7589-8877). **Author contributions:** Q.S. and Y.C. initiated the project. T.D., J.S., X.L. and Q.S. designed and performed the biochemical analysis. Y.C. and Z.X. performed the cryo-EM analysis. Q.S. and Y.C. analyzed the structure. X. S., X. L., and T. Z. purified Rabs. Z.C. and L.W. performed the AUC analysis. Q.S. and Y.C. wrote the manuscript. All the authors discussed the results and commented on the manuscript. Coordinates and structure factor of the structure reported here have been deposited into the Protein Data Bank with PDB Code: 6LT0, and EMDB code: EMD-0966.

## Abstract

A massive intronic hexanucleotide repeat (GGGGCC) expansion in *C9ORF72* is a genetic origin of amyotrophic lateral sclerosis (ALS) and frontotemporal dementia (FTD). Recently, C9ORF72, together with SMCR8 and WDR41, has been shown to regulate autophagy and function as Rab GEF. However, the precise function of C9ORF72 remains unclear. Here, we report the cryo-EM structure of the human C9ORF72-SMCR8-WDR41 complex at a resolution of 3.2 Å. The structure reveals the dimeric assembly of a heterotrimer of C9ORF72-SMCR8-WDR41. Notably, the C-terminal tail of C9ORF72 and the DENN domain of SMCR8 play critical roles in the dimerization of the two protomers of the C9ORF72-SMCR8-WDR41 complex. In the protomer, C9ORF72 and WDR41 are joined by SMCR8 without direct interaction. WDR41 binds to the DENN domain of SMCR8 by the C-terminal helix. Interestingly, the prominent structural feature of C9ORF72-SMCR8 resembles that of the FLNC-FNIP2 complex, the GTPase activating protein (GAP) of RagC/D. Structural comparison and sequence alignment revealed that Arg147 of SMCR8 is conserved and corresponds to the arginine finger of FLCN, and biochemical analysis indicated that the Arg147 of SMCR8 is critical to the stimulatory effect of the C9ORF72-SMCR8 complex on Rab8a and Rab11a. Our study not only illustrates the basis of C9ORF72-SMCR8-WDR41 complex assembly but also reveals the GAP activity of the C9ORF72-SMCR8 complex.

**Significance Statement:** C9ORF72, together with SMCR8 and WDR41, has been shown to form a stable complex that participates in the regulation of membrane trafficking. We report the cryo-EM structure of the C9ORF72-SMCR8-WDR41 complex at atomic resolution. Notably, the stoichiometry of the three subunits in the C9ORF72-SMCR8-WDR41 complex is 2:2:2. Interestingly, the C-termini of C9ORF72 and the DENN domain of SMCR8 mediate the dimerization of the two C9ORF72-SMCR8-WDR41 protomers in the complex. Moreover, WDR41 binds to the DENN domain of SMCR8 by the C-terminal helix without direct contact with C9ORF72. Most importantly, the C9ORF72-SMCR8 complex works as a GAP for Rab8a and Rab11a *in vitro,* and the Arg147 of SMCR8 is the arginine finger.

## Introduction

Amyotrophic lateral sclerosis (ALS) and frontotemporal dementia (FTD) are the most common neurodegenerative diseases and show overlapping pathology, genetic abnormalities, and patient symptoms(1–3). Disrupted RNA and protein homeostasis have been indicated to be the general causes of ALS-FTD(4). In 2011, the expanded intronic hexanucleotide repeat (GGGGCC) in the 5’ noncoding region of the gene *C9ORF72* was found to accounted for most cases of familial ALS and FTD as well as some sporadic cases of both diseases, representing a historic discovery(5–7). The two potential mechanisms of disease onset related to repeat expansion identified to date are gain-of-function and loss-of-function(3, 8). Studies suggest that neurotoxic materials, including dipeptide repeat proteins (DRPs) and RNA G-quadruplexes, are generated based on the expanded hexanucleotide repeat, which is described as “gain of function”(9–11). In addition, the expanded hexanucleotide repeat can also inhibit the transcription of *C9ORF72* and thereby decrease the production of C9ORF72 protein(5, 6, 12, 13). However, the precise function of C9ORF72 is unclear.

Recently, C9ORF72, together with SMCR8 (Smith-Magenis syndrome chromosomal region candidate gene 8) and WDR41 (WD40 repeat-containing protein 41), has been shown to form a stable complex that participates in regulating macroautophagy (hereafter referred to as autophagy) by directly interacting with the ULK1 complex(14–16). C9ORF72 knockout causes a defect in starvation-induced autophagy, indicating that C9ORF72 regulates autophagy positvely(14, 16–18). SMCR8 was so named because its gene location on the chromosome is close to the gene related to Smith-Magenis syndrome; however, it has no relationship with this syndrome(19). Interestingly, SMCR8 knockout results in an increase in ULK1 gene expression, which suggests that SMCR8 is a negative regulator of autophagy(20, 21). WDR41, the function of which is unknown, is a WD40 repeat-containing protein located on the ER (Fig. 1a)(22, 23). The subunits of the ULK1 complex, to which the C9ORF72-SMCR8-WDR41 complex (hereafter referred to as CSW) binds, are still controversial. Although studies have suggested that the CSW complex interacts with the ULK1 complex via ATG101 or FIP200(14, 16), the details of the interaction between these complexes have yet to elucidated.

**Figure 1.**
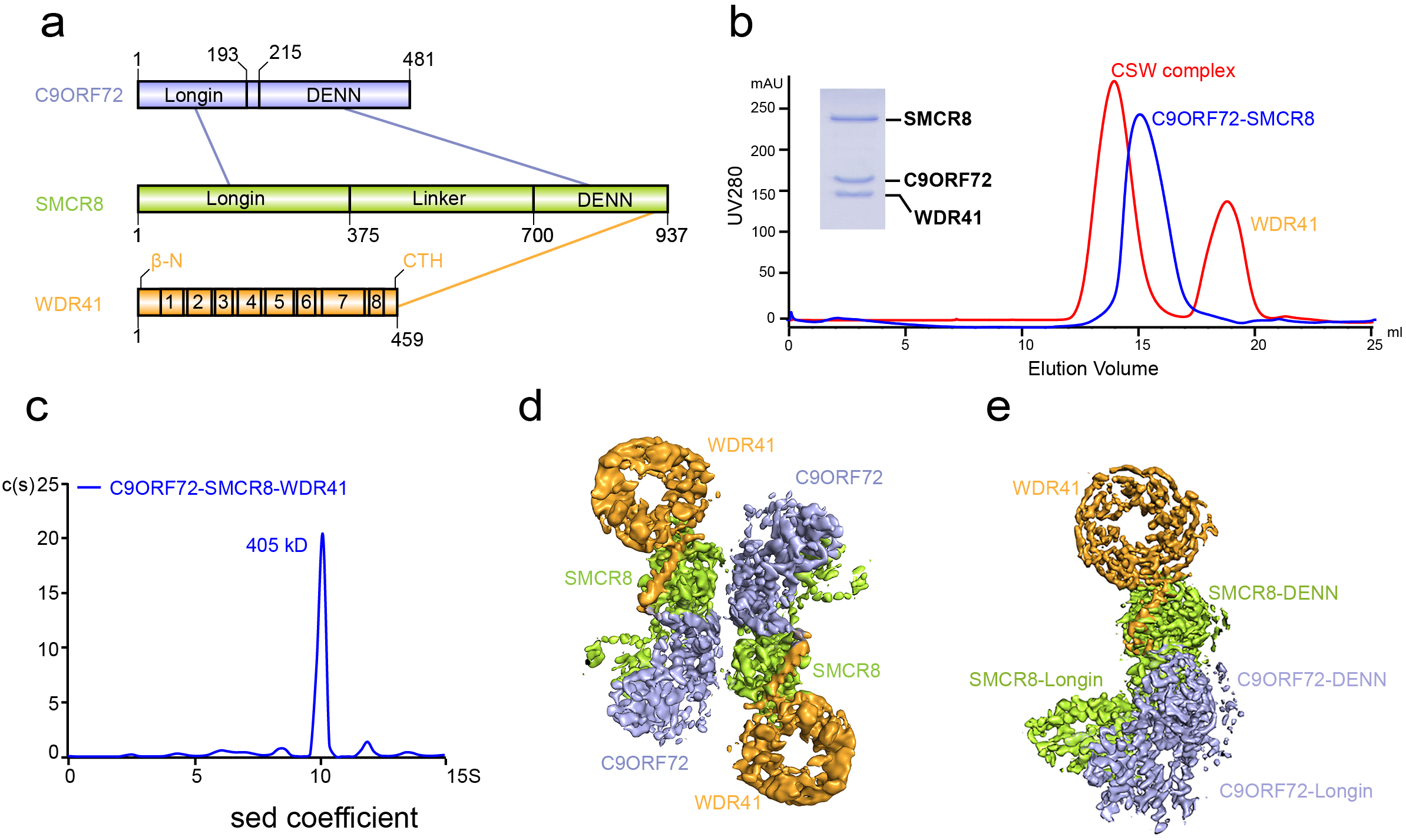
The CSW complex is a dimer of C9ORF72-SMCR8-WDR41. **A.** Schematic diagram of the domain arrangement of C9ORF72 (light-blue), SMCR8 (light-green), and WDR41 (orange). The names and boundaries of domains are labeled. β-N: N-terminal first β strand of WDR41. CTR: C-terminal helix of WDR41. The numbers in WDR41 represent WD40 domains: WD1 (41–81), WD2 (88–131), WD3 (137–168), WD4 (177–281), WD5 (226–276), WD6 (281–314), WD7 (326-401), and WD8 (411-432, and β-N). The interactions between different domains are shown in lines: light-blue line, C9ORF72-SMCR8 interaction; orange line, WDR41-SMCR8 interaction. **B.** Gel filtration (superpose 6 10/300 GL) profile of reconstituted C9ORF72-SMCR8-WDR41 and C9ORF72-SMCR8 complex. The horizontal axis is elution volume, and the vertical axis is UV absorption. The UV absorbance is shown in red (C9ORF72-SMCR8-WDR41) and blue (C9ORF72-SMCR8) lines. The peaks of proteins are labeled. The Coomassie blue-stained SDS-PAGE shows the peak fraction of the CSW from gel filtration. **C.** Analysis of peak fraction from (b) by Sedimentation velocity AUC. C(S) functions calculated from sedimentation velocity data are shown in blue curve. The calculated molecular mass is denoted. Horizontal axis: Sedimentation coefficient. Vertical axis: Continuous sedimentation coefficient distribution. (**D, E)** Cryo-EM density map of the CSW complex. **D**. The overall map for the dimer. **E.** The final map of one protomer of the CSW complex. The components of the CSW complex are indicated in different colors. Light-blue: C9ORF72; light-green: SMCR8; Orange: WDR41.

Both C9ORF72 and SMCR8 are predicted to be members of the DENN (differentially expressed in normal and neoplastic cells) family, although they share low sequence similarity (Fig. 1 a)(24–26). DENN domain-containing proteins are well known for functioning as guanine nucleotide exchange factors (GEFs) for many Rab GTPases(27–30). Hence, C9ORF72 and SMCR8 may participate in regulating membrane trafficking by mediating the activity of Rab GTPases(28)). Consistent with this hypothesis, studies showed that the CSW complex but not C9ORF72 alone has a significant ability to stimulate the exchange of GDP for GTP of Rab8a and Rab39b *in vitro*(14, 17).Besides, C9ORF72 has also been suggested to interact weakly with Rab11a(31). However, the physiological targets of the CSW complex need further analysis. Additionally, C9ORF72 and SMCR8 have been predicted to be similar to FLCN and FNIP. Before being shown to function as a GTPase activating protein (GAP) of RAG on lysosomes, the FLCN-FNIP complex was indicated to be a Rab35 GEF *in vitro(32–34).* Therefore, understanding the structure and biochemical properties of the CSW complex may shed light on the functions of this neurodegenerative disease-related complex.

In this manuscript, we determined the cryo-electron microscopy (cryo-EM) structure of the CSW complex, which revealed that the complex is composed of two copies of the three proteins, consistent with the biophysical analysis results. The C-terminal tail of C9ORF72 mediates the dimerization of two protomers of the CSW complex by binding to the DENN domain of SMCR8 in the other protomer, this observation was corroborated by biochemical analysis. In the protomer, C9ORF72 and WDR41 are held together by SMCR8 without direct contact with each other. WDR41 binds to the DENN domain of SMCR8 by the C-terminal helix. The overall structure of C9ORF72-SMCR8 resembles that of the FLNC-FNIP2 complex. Biochemical analysis revealed that C9ORF72-SMCR8 has GAP activity for Rab8a and Rab11a and that Arg147 of SMCR8 is critical to the GAP function. Together, our results reveal the assembly basis of the CSW complex and sheds light on the mechanism of its Rab GAP activity.

## Results

### The CSW complex is a dimer of trimer

The CSW complex was previously reported to form a stable complex at a stoichiometry of 1:1:1(17). After testing different expression systems, the well-behaved CSW complex was expressed and purified using Sf9 cells (Fig. 1b, S1). Curiously, the elution volume of the CSW complex in gel filtration corresponded to a molecular mass of ~400 kD, a mass larger than that of the CSW complex with a 1:1:1 stoichiometry of the three subunits (Fig. 1b, S1). To determine the accurate molecular mass of the CSW complex, an analytical ultracentrifugation (AUC) experiment was carried out. Interestingly, the measured sedimentation coefficient of the CSW complex was ~10S, which corresponded to a molecular mass of ~405 kD (Fig. 1c). The theoretical molecular masses of C9ORF72, SMCR8, and WDR41 are ~54 kD, ~105 kD, and ~48.5 kD, respectively. The intensity of Coomassie blue suggested that the molar ratio of the three subunits was 1:1:1 (Fig. 1b, S1). Collectively, these observations suggested that the CSW complex might contain two copies of each of the three subunits to yield a total molecular mass of 415 kD.

We carried out single-particle cryo-EM analysis of the CSW complex (Fig. S2, S3, Table S1) and solved the cryo-EM density map at a resolution of 3.2 Å (Fig. 1d, 1e). The resolution of the core domain ranges from 3.4 Å to 3.2Å, which clearly shows the side chains at the core region (Fig. S3), and provides us with the details of the CSW complex at atomic resolution. The map revealed that the CSW complex is a dimer of C9ORF72-SMCR8-WDR41, which is consistent with the data obtained from analytical gel filtration and AUC analysis (Fig. 1b, 1c, 1d). The model of the CSW complex was built by combining homology modelling and *de novo* model building, which allowed us to visualize the structure of the CSW complex in details.

### Both the C-terminal region of C9OFR72 and the DENN domain of SMCR8 are required for the dimerization of the CSW protomers

The CSW complex is a homodimer of C9ORF72-SWCR8-WDR41 with a width of ~ 130 Å and a height of 150 Å (Fig. 2a, S4). The two protomers present 2-fold symmetry and match well with each other after 180° rotation (Fig. 2a). The observed buried area between the two protomers is ~1544 Å^2^.

**Figure 2.**
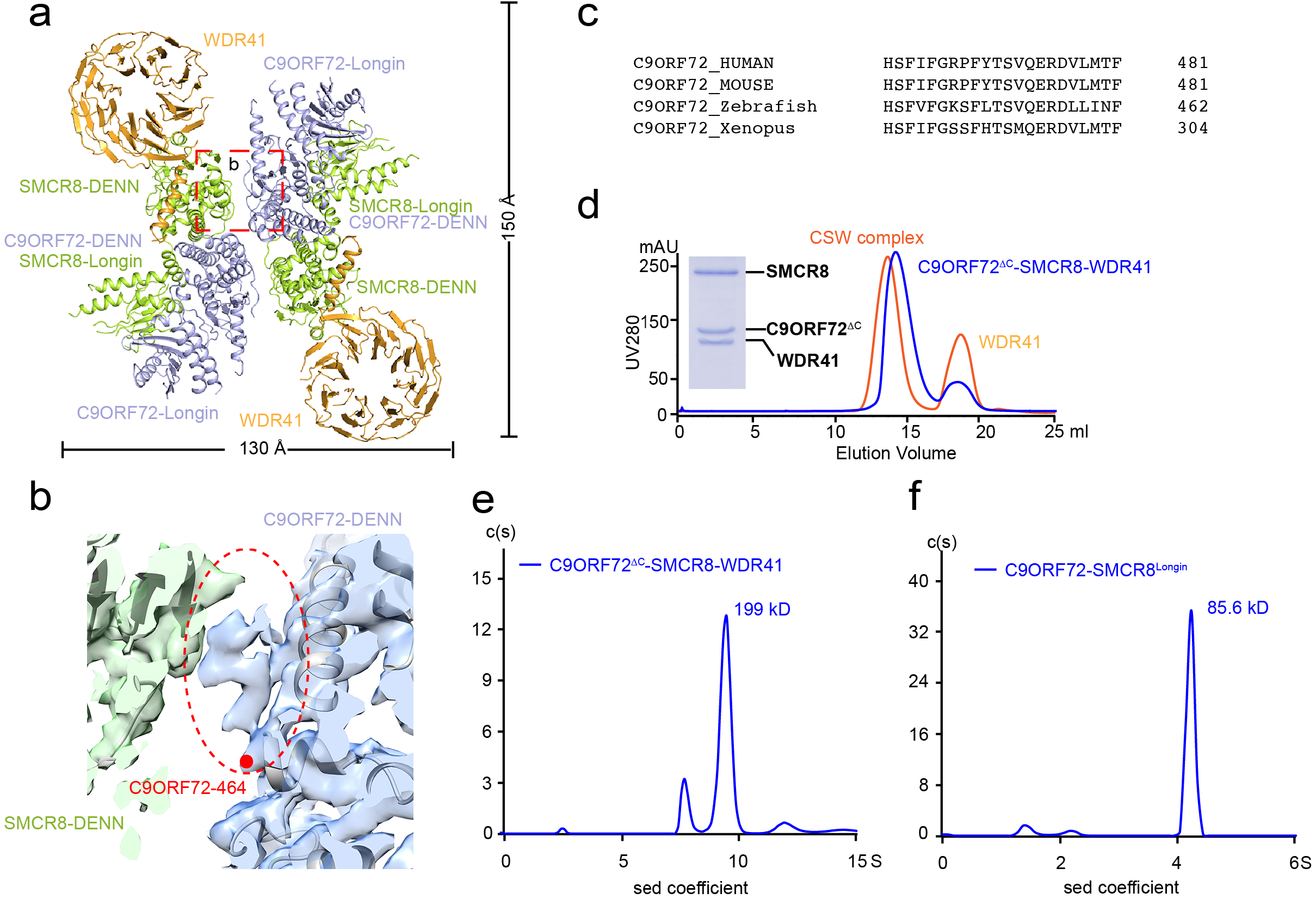
The dimer interface of the two protomers in the CSW complex. **A.** The overall structure of the CSW complex is shown in cartoon. The CSW complex shows a two-fold symmetry. The domains are labeled. The width and height are denoted. C9ORF72, SMCR8, and WDR41 are colored in light-blue, light-green, and orange, respectively. **B.** The density map of the interface between C9ORF72^CTR^ and SMCR8^DENN^. The unoccupied ambiguous map is highlighted with an oval circle. The last residue presented in the model was denoted with a red dot and labeled. **C.** The sequence alignment of C9ORF72^CTR^ in different species. HUMAN: *Homo sapiens;* Mouse: *Mus musculus;* Zebrafish: *Danio rerio;* Xenopus: *Xenopus tropicalis.* **D.** Gel filtration (superpose 6 10/300 GL) profile of reconstituted C9ORF72^ΔC^-SMCR8-WDR41 complex. C9ORF72^ΔC^ represents the construct of C9ORF72 with 461-481 deleted. The UV280 absorption curve of C9ORF72^ΔC^-SMCR8-WDR41 complex is shown in blue, whereas wild-type C9ORF72-SMCR8-WDR41 complex is shown in orange as comparison. The peak fraction of C9ORF72^ΔC^-SMCR8-WDR41 complex is examined by Coomassie blue stained SDS-PAGE. (**E, F**). Analysis of by Sedimentation velocity AUC of C9ORF72ΔC-SMCR8-WDR41 (**E**) and C9ORF72-SMCR8^Longin^ (**F**). C(S) functions calculated from sedimentation velocity data are shown in a blue curve. The calculated molecular mass is denoted. Horizontal axis: Sedimentation coefficient. Vertical axis: Continuous sedimentation coefficient distribution.

The structure model reveals that the dimerization of the two CSW protomers is mainly mediated by the C-terminal region of C9ORF72 and the DENN domain of SMCR8 (Fig. 2a, S4a). The density of the C9ORF72 C-terminal region (461-481, hereafter C9ORF72^CTR^) is ambiguous in both protomers (Fig. 2b). Moreover, C9ORF72^CTR^ is highly conserved cross species (Fig. 2c). To assess the contribution of C9ORF72^CTR^ to dimerization, a mutant with C9ORF72^CTR^ deleted (C9ORF72^ΔC^) was constructed. As expected, the C9ORF72^ΔC^-SMCR8 complex is capable of binding to WDR41 as tightly as the C9ORF72-SMCR8 complex (Fig. 2d, S1). However, the elution volume of the C9ORF72^ΔC^-SMCR8-WDR41 complex in gel filtration was delayed by ~0.9 ml relative to that of the CSW complex, which corresponded to a molecular mass of ~200 kD (Fig. 2d, S1). Consistent with this observation, the AUC experiment showed that the sedimentation coefficient of the C9ORF72^ΔC^-SMCR8-WDR41 complex is 9.87S, corresponding to a molecular mass of ~199 kD (Fig. 2e). Together, these observations indicate that the C9ORF72^ΔC^-SMCR8-WDR41 complex is monomeric in solution and that C9ORF72^CTR^ is critical to the dimerization of two CSW protomers.

The structure of CSW does not show where the C9ORF72^CTR^ packs on the DENN domain of SMCR8 (700-937, SMCR8^DENN^); therefore, the assembly status of the complex of C9ORF72 and SMCR8 Longin domain (1-349, SMCR8^Longin^) was evaluated. AUC analysis showed that the sedimentation coefficient of C9ORF72-SMCR8^Longin^ was ~4.35S (Fig. 2f), which corresponds to a molecular mass of ~ 85.6 kD and suggests C9ORF72-SMCR8^Longin^ is monomeric in solution. Gel filtration analysis indicated that C9ORF72-SMCR8^Longin^ is a monomer is solution as well (Fig. S5).

Collectively, the data support that both the C-terminal region of C9OFR72 and the DENN domain of SMCR8 together mediate the dimerization of the CSW protomers together.

### Organization of the CSW protomer

The C9ORF72-SMCR8-WDR41 complex in one protomer adopts an elongated rod shape, in which the C9ORF72-SMCR8 complex resembles the FLCN-FNIP2 complex (Fig. 3a)(35, 36). Structural comparison shows that C9ORF72 corresponds to FNIP2, whereas SMCR8 resembles FLCN (Fig. 3b, S6a), which is consistent with the bioinformatic analysis(24, 37).

**Figure 3.**
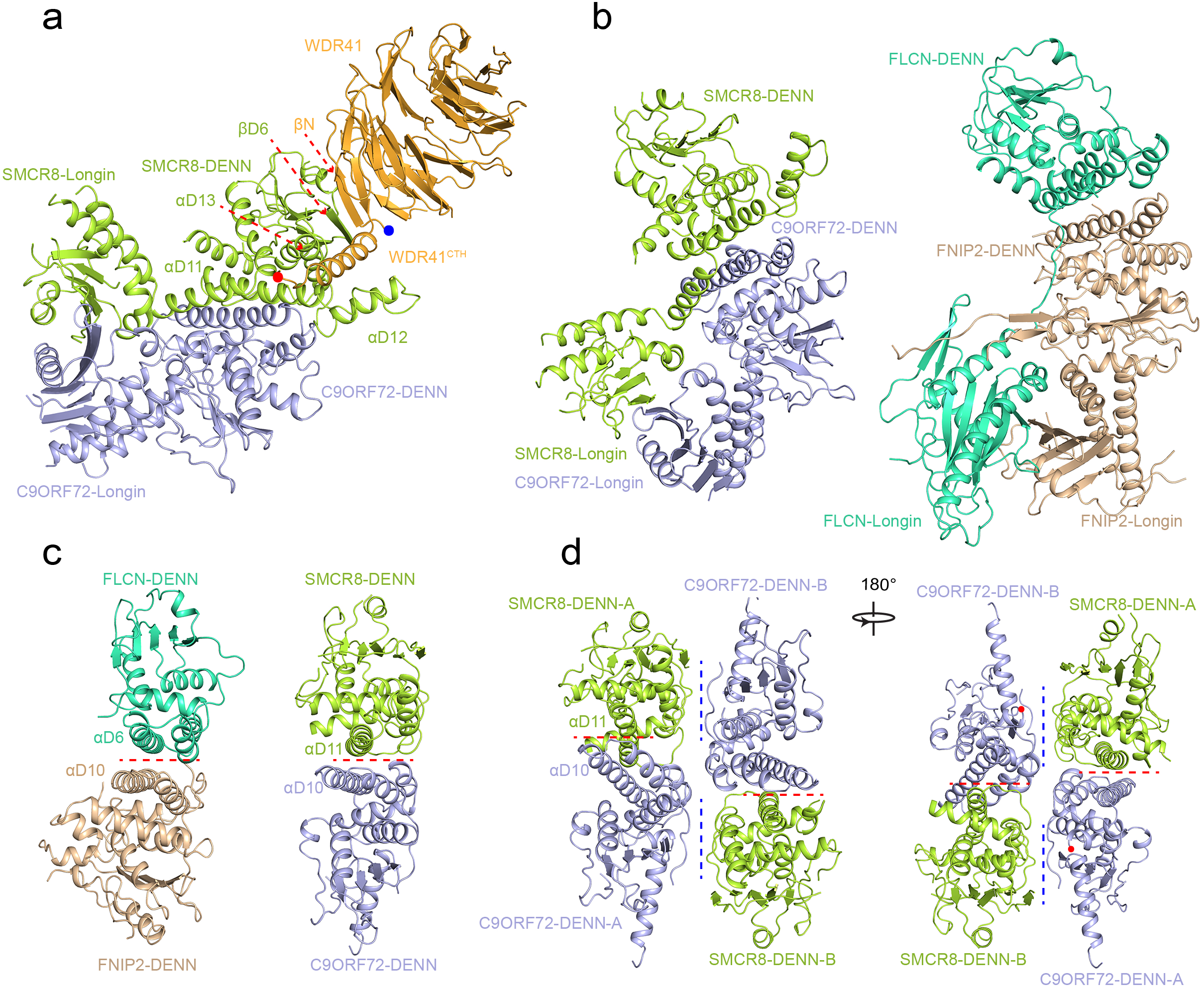
Organization of the CSW protomer. **A.** The overall structure of the CSW protomer. The key secondary structures are labeled. The colors are consistent with the previous figures **B.** The structure of the C9ORF72-SMCR8 complex adopts a similar organization to the FLCN-FNIP2 complex. Left: C9ORF72-SMCR8 complex. Right: FLCN-FNIP2 complex. C9ORF72, SMCR8, and WDR41 are colored as (**A**). FLCN and FNIP2 are colored in greencyan and wheat. The details of the comparison are presented in Fig. S5. (**C, D**) Two interfaces between C9ORF72^DENN^ and SMCR8^DENN^ are shown. **C.** Comparison of DENN pair of C9ORF72-SMCR8 and FLCN-FNIP. The key helixes are labeled. The red dash line indicates the position of the interface. **D.** The two interfaces between C9ORF72^DENN^ and SMCR8^DENN^ are shown. The red dash line indicates the positions of the intra-protomer interface, while the blue dash line indicates the positions of the interprotomer interface.

Superimposition of the C9ORF72-SMCR8 complex and FLCN-FLIP2 dimer revealed that the position of the Longin domain shifts by approximately 40 Å when the DENN domains are superimposed well (Fig. S6a). Although the C9ORF72-SMCR8 complex and FLCN-FNIP2 dimer show a slight difference in overall conformation, the Longin and DENN dimer of the C9ORF72-SMCR8 complex can be superimposed well on their counterparts in FLCN-FNIP2 (Fig. 3b, S6b, c). Intriguingly, SMCR8^Longin^ and SMCR8^DENN^ are connected by αL5, αL6 and the following loop, whereas the Longin domain and DENN domain of FLCN are connected by a flexible loop (Fig. S4b). This observation might explain the structural discrepancy between the two complexes.

The interaction of C9ORF72 and SMCR8 in one protomer is mediated by the Longin domains and DENN domains, respectively, which is consistent with the observations in the FLCN-FNIP2 structure (Fig. 3b, c, S6a, b, c)(35, 36). Importantly, C9ORF72^CTR^ and SMCR8^DENN^ also mediate the dimerization of the CSW protomers (Fig. 2a, b). Thus, there are two interfaces between the DENN domains of C9ORF72 and SMCR8: the interface consisting of αD10 of C9ORF72 and αD11 of SMCR8, which mediates the intra-protomer interaction, and the interface between C9ORF72^CTR^ and SMCR8^DENN^ mediates the dimerization of the CSW protomer (Fig. 3d). This feature distinguished the DENN domains of the C9ORF72-SMCR8 protomer from other members of the DENN domain-containing proteins.

The structure of WDR41 reveals an 8-blade β-propeller with the N-terminal first strand and C-terminal last three strands coming together to form the first propeller, generating a “Velcro” closure that stabilizes the architecture of WDR41 (Fig. 3a, S4d)(22). WDR41 binds to the SMCR8 DENN domain via its N-terminal β-strand (βN) and C-terminal helix (CTH) without direct physical contact with C9ORF72 (Fig. 3a, S4a, S4d).

### The interface of SMCR8-WDR41

As mentioned above, WDR41 binds to SMCR8 mainly via its C-terminal helix (WDR41^CTH^) and the very N-terminal β-strand (Fig. 3a). WDR41^CTH^ packs against the groove formed by αD11, αD12, and αD13 of SMCR8, whereas the N-terminal β-strand of WDR41 forms an antiparallel β-sheet with βD6 of SMCR8 (Fig. 3a, 4a).

**Figure 4.**
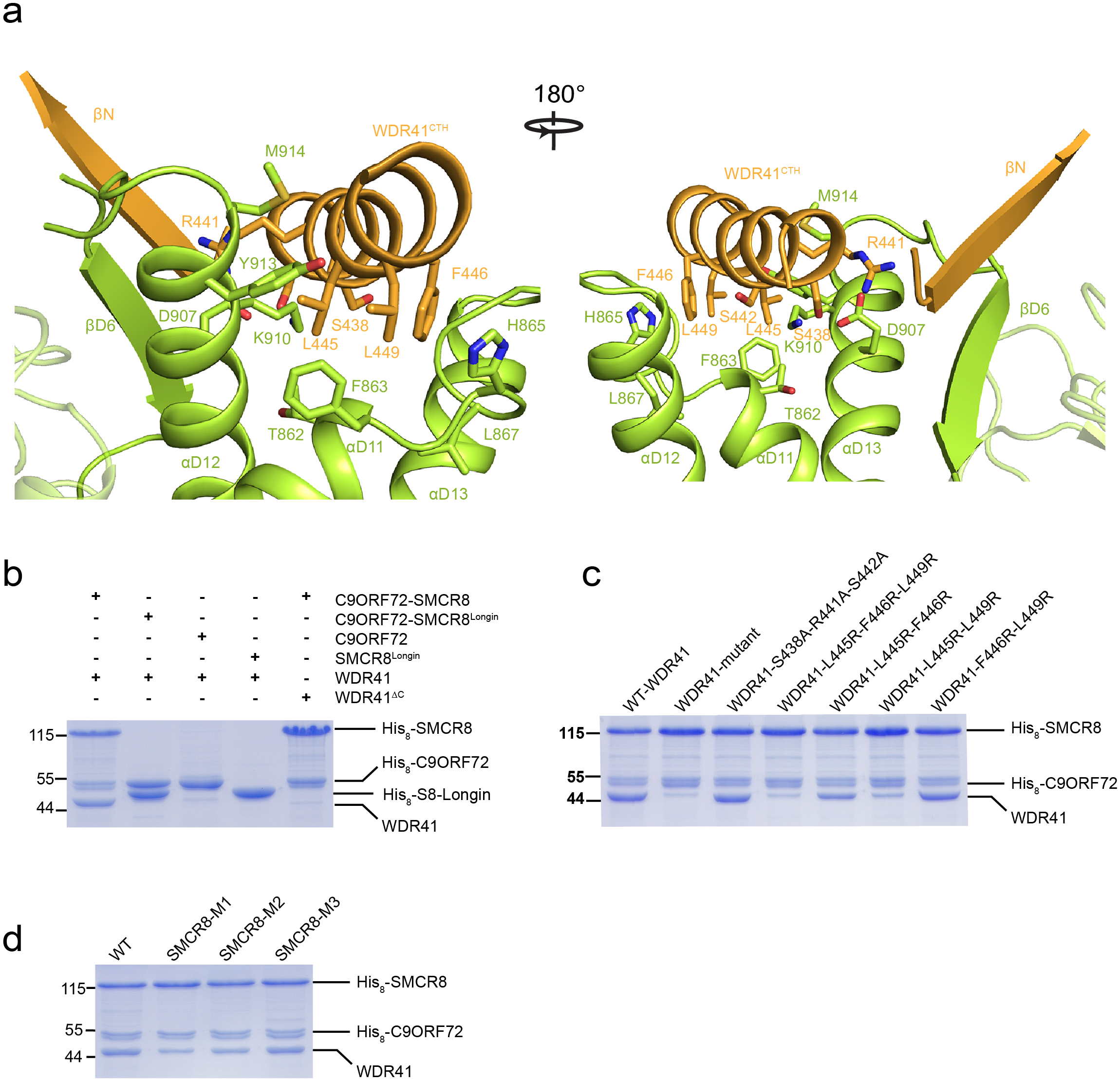
The interface of SMCR8 and WDR41. **A.** The details of the interface of SMCR8 and WDR41. The key residues on the interface are shown in stick model. Secondary structures are shown as cartoon. Left: front view; right: back view. **B.** C9ORF72, SMCR8^Longin^, C9ORF72-SMCR8^Longin^ complex, and C9ORF72-SMCR8 complex were used to pulldown WDR41 and WDR41^ΔC^. **(C, D)** Mutational analysis of key residues on WDR41^CTH^ (**C**) and SMCR8 (**D**) using pulldown assay. The results are visualized by Coomassie blue-stained SDS-PAGE. Protein markers are labeled at left (unit: kD). **c.** Wild-type C9ORF72-SMCR8 complex with N-terminal His-tag was used to pull down untagged wild-type or indicated mutants of WDR41. WDR41-mutant: WDR41 with mutations of S438A-R411A-S442A-L445R-F446R-L449R. Other mutants are shown as the label. **D.** Indicated mutants of SMCR8 with N-terminal His-tag were used to pull down untagged wild-type WDR41. WT: wild-type C9ORF72-SMCR8; SMCR8^M1^: C9ORF72-SMCR8^M1^ (SMCR8 with mutations of T862A, F863A, H865A, L867A, E907A, K910A, Y913A, M914A); SMCR8^M2^: C9ORF72-SMCR8^M2^(SMCR8 with mutations of T862A, F863A, H865A, L867A); SMCR8^M3^: C9ORF72-SMCR8^M3^ (SMCR8 with mutations of E907A, K910A, Y913A, M914A).

To verify this observation, a mutant of WDR41 with a C-terminal helix deleted (436-459, WDR41^ΔC^) and SMCR8^Longin^ were purified successfully and subjected to the pulldown assay. In the pulldown assay, C9ORF72-SMCR8 was able to interact with WDR41 but not WDR41^ΔC^ (Fig. 4b, S7a). In contrast, C9ORF72 was not capable of binding to WDR41 or WDR41^ΔC^ (Fig. 4b, S7a). Moreover, SMCR8^Longin^ was not able to bind to either WDR41 or WDR41^ΔC^ (Fig. 4b, S7a). Collectively, these observations support that WDR41 binds to the DENN domain of SMCR8 via its C-terminal helix.

The density map of the interface was clear enough to carry out a detailed alanine mutagenesis analysis. The residues on WDR41^CTH^ facing SMCR8, including S438, R441, S442, L445, F446, and L449, were individually mutated to either alanine or arginine (Fig. S7b). However, none of the mutated residues disrupted the interaction between WDR41 and the C9ORF72-SMCR8 complex (Fig. S7b). Therefore, combinations of these mutations were examined. The mutant (WDR41-mutant) with all six key residues mutated abolished the interaction of WDR41 and SMCR8 (Fig. 4c, S7c). Interestingly, WDR41^L445R-F446R-L449R^ blocked binding to the C9ORF72-SMCR8 complex, whereas WDR41^S438A-R441A-S442A^ had little effect on binding (Fig. 4c, S7c). The double mutant WDR41^L445R-L449R^ had obvious effects on binding, whereas WDR41^L445R-F446R^ and WDR41^F446R-L449R^ had weaker phenotypic effects than WDR41 ^L445R-F446R-L449R^ (Fig. 4c, S7c).

On the SMCR8 side, all residues (T862-F863-H865-L867-E907A-K910-Y913-M914) involved in binding to L445, F446, and L449 of WDR41 were mutated to alanine, and the resulting construct was termed SMCR8^M1^. The C9ORF72-SMCR8^M1^ complex was purified and subjected to a pulldown assay. Compared with wild-type C9ORF72-SMCR8, C9ORF72-SMCR8^M1^ reduced the binding ability to WDR41 by ~50% (Fig. 4d, S7d). Two other mutants, SMCR8^M2^ (T862A-F863A-H865A-L867A) and SMCR8^M3^ (E907A-K910A-Y913A-M914A), were also tested and showed little effect on the binding to WDR41 (Fig. 4d, S7d). The alanine mutational analysis was probably unable to completely destroy the hydrophobic cavity. Hence, we attempted to mutate these residues to arginine, however, a single mutation of any residues to arginine caused C9ORF72-SMCR8 to precipitate. We also tested the effect of βD6 of SMCR8 on the binding to WDR41 by deleting the C-terminal region of SMCR8 (923-937, SMCR8^ΔC^). The pulldown assay showed that SMCR8^ΔC^ had little effect on the interaction of C9ORF72-SMCR8 and WDR41 (Fig. S7e), indicating that the antiparallel β-sheet formed between βD6 of SMCR8 and βN of WDR41 is not essential to the binding of SMCR8 and WDR41.

Collectively, these observations demonstrate that the hydrophobic interaction of the C-terminal helix of WDR41 and the DENN domain of SMCR8 is critical to the binding of WDR41 to SMCR8.

### The C9ORF72-SMCR8 complex is a GAP for Rab8a *in vitro*

The CSW complex has been reported to function as a GEF of Rab7a, Rab8a, Rab11a, Rab39a, and Rab39b(14, 17, 31). Hence, a bioluminescence-based GTPase activity assay was carried out to assess whether the CSW complex stimulates these Rabs(38). In this type of assay, unhydrolyzed GTP can be transformed into a bioluminescence signal. A higher the bioluminescence signal indicates less GTP hydrolyzation by Rabs. Stimulated by either the CSW complex or the C9ORF72-SMCR8 complex but not C9ORF72 alone, the amount of GTP consumed by Rab8a and Rabi 1 a increased by approximately 100% and 50%, respectively (Fig. 5a). However, C9ORF72, the CSW complex, and the C9ORF72-SMCR8 complex had little effect on Rab7a, Rab39a, and Rab39b (Fig. 5a). Interestingly, the CSW complex and C9ORF72-SMCR8 complex showed similar effects on Rab8a and Rab11a, indicating that WDR41 is not essential for the stimulation of target Rabs (Fig. 5a). Moreover, Rab8a and Rab11a play critical roles in coordinating primary ciliogenesis and axon growth(39, 40). Therefore, we focused on the relationships between Rab8a/11a and the CSW complex and the C9ORF72-SMCR8 complexes *in vitro*.

**Figure 5.**
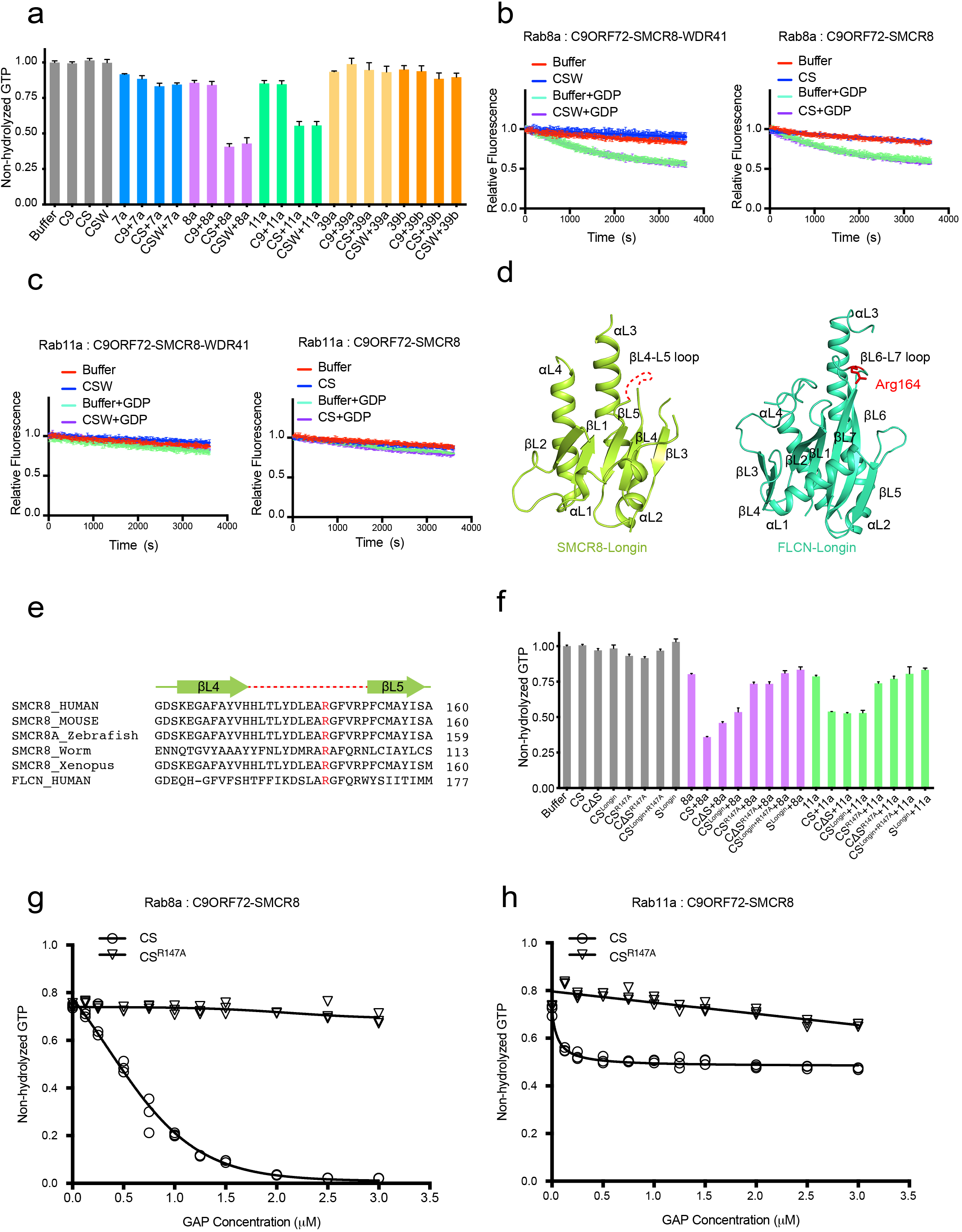
CSW and C9ORF72-SMCR8 complex are GAPs of Rab8a and Rab11a *in vitro.* **A.** The screen of Rabs that can be stimulated by CSW or C9ORF72-SMCR8 complex using bioluminescence based GTPase activity assay. In this assay, the final concentration of Rabs and C9/CS/CSW was 1.5 μM and 0.75 μM, respectively. The GTP in the reaction system containing no protein was normalized to “1.0”. The proteins or protein mixtures added in the reaction system are indicated below. Buffer: Buffer control, C9: C9ORF72, CS: C9ORF72-SMCR8, CSW: wild-type C9ORF72-SMCR8-WDR41, 7a: Rab7a, 8a: Rab8a, 11a: Rab11a, 39a: Rab39a, 39b: Rab39b. The error bars represent mean ± SD (N=3). (**B, C**) mant-GDP based nucleotide exchange assay for Rab8a (**B**) and Rab11a (**C**). The abbreviations of protein names are same to the panel (**A**). 100 μM GDP was used to initiate the exchange reaction. Reactions without GDP were monitored as a control. The error bars represent mean ± SD (N=3). The intrinsic nucleotide exchange rate of Rab11a is negligible. **D**. Comparison of the Longin domains of SMCR8 and FLCN. The secondary structures are labeled. The Arg164 of FLCN-Longin domain is highlighted in red. The missing loop of SMCR8^Longin^ is indicated as a red dash line. **E**. Sequence alignment of the βL4-βL5 loop of SMCR8. Arg147 of human SMCR8 is conserved cross species and corresponds to Arg164 of FLCN. HUMAN: *Homo sapiens;* Mouse: *Mus musculus;* Zebrafish: *Danio rerio;* Worm: *Caenorhabditis elegans;* Xenopus: *Xenopus tropicalis;* The secondary structures are shown on top of the sequence in light-green, and the βL4-βL5 is shown in red dash line. **F**. Test the effect of Arg147 of SMCR8 on the stimulation of Rab8a and Rab11a. The experiment was carried out as panel (**A**), and the labels are same to the panel (**A**).CΔS: C9ORF72^ΔC^-SMCR8, CS^Longin^: C9ORF72-SMCR8^Longin^, CS^R147A^: C9ORF72-SMCR8^R147A^, CΔS^R147A^: C9ORF72^ΔC^-SMCR8^R147A^, CS^Longin+R147A^: C9ORF72-SMCR8^Longin+R147A^, S^Longin^: SMCR8^Longin^. The error bars represent mean ± SD (N=3). (**G, H**) Measurement of GAP activity to Rab8a (**G**) and Rab11a (**H**) using different concentrations of C9ORF72-SMCR8 or C9ORF72-SMCR8^Arg147^. The final concentration of Rabs in this assay was 2 μM. The concentrations of C9ORF72-SMCR8 or C9ORF72-SMCR8^Arg147^ are indicated in the horizontal axis. The relative amount of non-hydrolyzed GTP in the system is shown in the vertical axis. Each concentration of GAP was measured in triplicate. The value of each measurement is shown in as “◦” (C9ORF72-SMCR8) and “▽” (C9ORF72-SMCR8^Arg147^) since the error bars of several measurements are too small to be shown clearly in the figure. The curve was fitted to using the Stimulation Model in Graphpad.

Since the CSW complex was previously identified as Rab GEF, we investigated whether the CSW complex and C9ORF72-SMCR8 complex promote the nucleotide exchange rate of Rab8a and Rab11a. Hence, a fluorescence-based GEF activity assay was performed using N-methylanthraniloyl (MANT)-GDP(41, 42). Compared with buffer, the CSW complex and the C9ORF72-SMCR8 complex were unable to accelerate mant-GDP release from Rab8a and Rab11a (Fig. 5b, c), suggesting that neither complex functions as a GEF for Rab8a and Rab11a.

As previously mentioned, the C9ORF72-SMCR8 complex shares a similar overall structure with the FLCN-FNIP2 heterodimer, the GAP of RagC/D(33–36). Arg164 on the loop of β6-β7 of FLCN is critical to the GAP activity of the FLCN-FNIP2 complex(35, 36). Although the density of the β6-β7 loop in the C9ORF72-SMCR8 complex is not clear, sequence alignment shows that Arg147 of SMRC8 is conserved across species and corresponds to Arg164 of FLCN (Fig. 5d, e). Hence, we investigated whether the C9ORF72-SMCR8 complex has GAP activity for Rab8a and Rab11a. The effect of Arg147 of SMCR8 was probed by the mutation R147A using a bioluminescence assay.

Excitingly, compared with wild-type C9ORF72-SMCR8, C9ORF72-SMCR8^R147A^ showed no obvious stimulatory effect on Rab8a and Rab11a (Fig. 5f). Intriguingly, C9ORF72^ΔC^-SMCR8 showed a stimulatory effect on Rab8a and Rab11a similar to that of wild-type C9ORF72-SMCR8, suggesting that the dimerization of CSW protomers is not essential for the stimulatory effect on Rab8a and Rab11a (Fig. 5f). Moreover, C9ORF72^ΔC^-SMCR8^R147A^ eliminated the stimulatory effect on Rab8a and Rab11a (Fig. 5f).

C9ORF72 showed no stimulatory activity against Rab8a and Rab11a. Hence, we attempted to test whether SMCR8 alone had GAP activity against Rab8a and Rab11a. However, SMCR8 cannot be purified alone. Fortunately, we were able to successfully purify SMCR8^Longin^ and C9ORF72-SMCR8^Longin^. Interestingly, SMCR8^Longin^ showed no stimulatory effects on Rab8a and Rab11a (Fig. 5f), whereas C9ORF72-SMCR8^Longin^ showed an effect on Rab8a and Rab11a similar to that of C9ORF72-SMCR8 (Fig. 5f). As expected, C9ORF72-SMCR8^Longin-R147A^ showed no GAP stimulatory effects on Rab8a and Rab11a (Fig. 5f).

Together, the data indicate that CSW and the C9ORF72-SMCR8 complex but not C9ORF72 or SMCR8 alone can exhibit GAP activity for Rab8a and Rab11a *in vitro.* Additionally, the data support that the dimerization of C9ORF72-SMCR8 is not required for the complex’s GAP activity.

To further assess the responses of Rab8a and Rab11a to stimulation by the C9ORF72-SMCR8 complex, a titration experiment was performed at a fixed concentration of Rab. With increasing concentration of the wild-type C9ORF72-SMCR8 complex, Rab8a hydrolyzed more GTP and almost exhausted the GTP when the molar ratio of Rab8a and C9ORF72-SMCR8 complex reached 1:1. In contrast, the C9ORF72-SMCR8^R147A^ complex showed no obvious stimulatory effect on Rab8a (Fig. 5g).

Similarly, Rab11a consumed more GTP with increasing concentration of the C9ORF72-SMCR8 complex but not that of the C9ORF72-SMCR8^R147A^ complex (Fig. 5h). However, the limit of GTP hydrolyzed by stimulated Rab11a was ~ 50% of the total amount of GTP (Fig. 5h). This phenomenon might have been caused by the slow intrinsic nucleotide exchange rate of Rab11a (Fig. 5c), which might have prevented Rabi i a from efficiently accessing GTP when the ratio between GTP:GDP in the system was 1:1(43).

In a word, C9ORF72-SMCR8 showed GAP activity against Rab8a and Rab11a *in vitro,* which was contrary to our expectation. Furthermore, Arg147 of SMCR8 is critical to the GAP activity of C9ORF72-SMCR8.

## Discussion

The CSW complex is critical to a variety of cellular processes and is strongly associated with familial ALS and FTD. In addition, the CSW complex is highly conserved in animals (Fig. S9–11). Nevertheless, the exact functions of the CSW complex remain unclear. Here, we investigated this important complex using recombinant proteins, biochemical assays, and cryo-EM analysis.

First, we determined that the stoichiometry of C9ORF72, SMCR8, and WDR41 in the CSW complex is 2:2:2. Both biochemical analysis and the structure of the CSW complex revealed that the CSW complex is a dimer of C9ORF72-SMCR8-WDR41. Although the density map of the dimerization interface is unclear, mutagenesis analysis, AUC analysis and analytical gel filtration indicated that C9ORF72^CTR^ and SMCR8^DENN^ are necessary for the dimerization of C9ORF72-SMCR8-WDR41. Notably, dimerization of C9ORF72-SMCR8-WDR41 or C9ORF72-SMCR8 is not essential to the complex’s GAP activity (Fig. 5f). To obtain detailed information on the dimer interface of C9ORF72-SMCR8-WDR41, a higher-resolution structure is needed, and determining the biological significance of dimerization requires *in vivo* genetic and functional analyses.

Additionally, we clarified the relationship among C9ORF72, SMCR8, and WDR41 based on the structure of CSW. WDR41 binds to a groove on SMCR8^DENN^ without direct contact with C9ORF72 (Fig. 4). WDR41 and C9ORF72 are joined together by SMCR8. A triad of L445, F446, and L449 from WDR41^CTH^ is critical to the binding of WDR41 to SMCR8. Mutants containing L445R, F446R, and L449R do not have the ability to bind to SMCR8. Eight residues of SMCR8^DENN^ involved in the binding to WDR41^CTH^ were assessed by alanine mutagenesis analysis since arginine mutagenesis crashed SMCR8. All eight residues have to be mutated into alanine simultaneously to cripple the binding of SMCR8 to WDR41. Interestingly, WDR41 is not essential to the GAP activity of the CSW complex. Previous studies showed that WDR41 locates on ER and participates in the autophagosome-lysosome pathway(15, 16, 23). Thus, the role of WDR41 in the CSW complex might be localizing the complex on appropriate positions.

We found that the DENN domain of C9ORF72 can interact with two DENN domains of SMCR8 simultaneously, *vice versa* (Fig. 3d). Not only does such a binding mode enable the DENN dimer of C9ORF72-SMCR8 to mediate intra-protomer interaction as well as the inter-protomer interaction, it distinguishes these DENN domains from others, including the DENN domains of FLCN and FNIP2. These observations enhance our understanding of the functional evolution of the DENN domain.

Most importantly, the GAP activity of C9ORF72-SMCR8 was identified based on the structure of C9ORF72-SMCR8 and a GTPase activity assay. The structure of C9ORF72-SMCR8 resembles that of FLCN-FNIP2, which is a GAP of RagC/D. The Longin dimers in both structures adopt similar organization patterns (Fig. S4c). Intriguingly, Lst7, the homologue of FLCN in yeast, contains only the Longin domain(44, 45). Moreover, the catalytic arginine of FLCN-FNIP2 is located in the Longin domain of FLCN(35, 36). Together, these data indicate that the Longin dimer might be functionally conserved during evolution. Based on the structure and sequence alignment, Arg147 of SMCR8 was found to correspond to Arg164. However, previous studies indicate that the CSW complex might function as a GEF of many Rabs. Hence, we screened Rabs that can be stimulated by CSW or the C9ORF72-SMCR8 complex *in vitro* using reconstituted proteins. Serendipitously, CSW and C9ORF72-SMCR8 were found able to stimulate Rab8a and Rab11a at similar levels, which indicated that WDR41 is not essential for the stimulating effect of CSW on Rab8a and Rab11a. Curiously, neither CSW nor C9ORF72-SMCR8 accelerated the nucleotide exchange rate of Rab8a and Rab11a *in vitro,* indicating that CSW and C9ORF72-SMCR8 are not GEFs for Rab8a and Rab11a. Mutagenesis analysis combined with GAP-stimulated GTPase activity assay confirmed that Arg147 of SMCR8 is essential to its GAP activity for Rab8a and Rab11a *in vitro*. In addition, both C9ORF72 and SMCR8 are required for GAP activity, as evidenced by the finding that neither C9ORF72 nor SMCR8^Longin^ shows GAP activity toward Rab8a and Rab11a. It is of interest to dissect how C9ORF72 and SMCR8 recognize and collaboratively activate Rab8a/11 a. Unfortunately, we have not yet been able to obtain a stable complex of C9ORF72-SMCR8-Rab8a/11a, indicating that the interaction of C9ORF72-SMCR8 and Rab8a/11a might be weak or ephemeral.

Our results demonstrate that CSW is a dimer of C9ORF72-SMCR8-WDR41 and is able to stimulate the GTPase activity of Rab8a and Rab11a as a GAP. Rab8a and Rab11a function coordinately in many fundamental cellular processes, including receptor recycling, ciliogenesis, and axon growth(39, 40, 46). It is tempting to propose the following model: CSW takes two endosomes containing Rab8a/11a via WDR41 simultaneously and then activates the Rabs located on the surface to prompt the fusion of the two endosomes.

Although the GAP activity of the C9ORF72-SMCR8 complex has been tested *in vitro*, the specific biological processes in which CSW and the C9ORF72-SMCR8 complex participate as a GAP require further exploration. The findings and the biochemical data presented in our manuscript will definitely facilitate future functional studies of CSW and the C9ORF72-SMCR8 complex.

## Materials and methods

### protein purification

Protein were expressed in *E. coli* BL21 (DE3) cells or Sf9 cells and purified using FPLC chromatography.

### Analytical ultracentrifugation

Analytical Ultracentrifugation Sedimentation experiments were carried out at 20 °C in an XL-I analytical ultracentrifuge (Beckman-Coulter) equipped with Rayleigh Interference detection (655 nm).

### Structure determination

The structure of CSW complex were determined using cryo-EM.

### GTPase activity assay

GTPase activity assays were carried out using GTPase-Glo assay kit (Promega, V7681).

### Nucleotide exchange assay

Releasing of MANT-GDP was recorded by monitoring the decrease in fluorescence emission at 448 nm (excited at 360 nm) in intervals of 15 s at 25 °C for 3600 seconds. Data were collected using 384-well plate (Corning, 3701) in BioTek Synergy2.

## Acknowledgements

We thank the Tsinghua University Branch of China National Center for Protein Sciences (Beijing) for providing the cryo-EM facility support. We thank the computational facility support on the cluster of Bio-Computing Platform (Tsinghua University Branch of China National Center for Protein Sciences Beijing). We thank Jianhua He, Bo Sun, Wenming Qin, Huan Zhou, Feng Yu from BL17U, BL18U, and BL19U at National Facility for Protein Science in Shanghai, Zhangjiang Laboratory (NFPS, ZJLab), China for providing technical support and assistance in crystal testing. We also thank staff from Protein Preparation and Identification Facility at Technology Center for Protein Science, Tsinghua University for the assistance with AUC data collection. This work was supported by National Key R&D Program of China grant 2017YFA0506300 (Q.S.), 2018YFC1004601 (S.Q.), and NSFC grants 81671388 (Q. S.), 31770820 (L. K.).

## Materials and methods

### Cloning and protein purification

The DNAs of human C9ORF72 and SMCR8 were subcloned into pFastBac dual under control of the polyhedron promoter and the p10 promoter, respectively. The DNA of WDR41 was subcloned into pFastBac dual under control of the polyhedron promoter with a N-terminal His_8_-tag that can be removed by Tobacco Etch Virus (TEV) protease. Both C9ORF72 and SMCR8 were expressed with N-terminal His8-tag that can be removed by TEV. The baculoviruses were generated in Sf9 cells with the bac-to-bac system (Life Technologies). The cells were infected and harvested after 60 hours. Cells were pelleted by centrifugation at 2000 x g for 15 minutes. The pellets were lysed in 25 mM Tris-HCl pH 8.0, 150 mM NaCl, 0.5 mM TCEP-HCl, 1 mM PMSF, 0.8 μM Aprotinin, 1 μM Pepstatin, and 10 μM Leupeptin by French Press. The lysate was then centrifuged at 25,000 x g for 30 min at 4 °C. The supernatants of C9ORF72-SMCR8 complex and WDR41 were loaded onto Ni-NTA resin at 4 °C, respectively. C9ORF72-SMCR8 complex and WDR41 were eluted with 25 mM Tris-HCl pH 8.0, 150 mM NaCl, 0.5 mM TCEP-HCl, and 250 mM Imidazole pH 8.0, respectively. The eluate of target proteins was diluted 5-fold with 25 mM Tris-HCl pH 8.0, 2 mM DTT and applied to a Hi-Trap Q HP column. Peak fractions were pooled and digested with TEV protease at 4 °C overnight. TEV and His-tag were removed by loading the solution onto Ni-NTA resin. Target proteins were further purified on a Superose 6 10/300 GL column equilibrated with 25 mM Tris-HCl pH 8.0, 150 mM NaCl, and 2 mM DTT. The peak fractions were pooled and flash-frozen in liquid N^2^ for storage.

To reconstitute the CSW complex, purified C9ORF72-SMCR8 complex and WDR41 were mixed at a molar ratio of 1:2 and incubated at 4 °C for 45 minutes. Then the mixture was applied to Superose 6 10/300 column equilibrated with 25 mM Tris-HCl pH 8.0, 150 mM NaCl, and 2 mM DTT (or 0.5mM TCEP). Peak fractions were identified using SDS-PAGE and flash-frozen in liquid N2 for storage.

SMCR8^Longin^ was expressed in *E. coli* BL21 (DE3) cells with N-terminal His6-tag that can be removed by TEV. After induction with 0.2 mM IPTG overnight at 16 °C, the bacteria were pelleted by centrifugation at 4000 x g for 10 minutes. Target proteins were purified as described above.

C9ORF72 and the C9ORF72-SMCR8^Longin^ complex was expressed in Sf9 cells and purified as the C9ORF72-SMCR8 complex.

The primers for point mutations were designed facilitated by a python script(1). All the mutant constructs were verified by DNA sequencing, and all the mutants were purified as the wild type proteins.

### Analytical ultracentrifugation

Analytical Ultracentrifugation Sedimentation experiments were carried out at 20 °C in an XL-I analytical ultracentrifuge (Beckman-Coulter) equipped with Rayleigh Interference detection (655 nm). BTN3A1 B30.2 (400 mL, 27 mM, 54 mM, 160 mM, 240 mM respectively) were centrifuged at 50,000 or 30,000 rpm for 8 hours, in an An50Ti rotorusing12 mm double-sector aluminum center pieces. All samples were prepared in the interaction buffer (25 mM Tris pH 8.0, 150 mM NaCl, 0.5 mM TCEP). Interference profiles were recorded every 6 min. Data analysis was conducted with the software Sedfit 11.

### EM sample preparation and data collection

The purified CSW complex was concentrated to approximately 1 mg/mL. Aliquots (4 μL) of the protein complex were placed on glow-discharged holey carbon grids (Quantifoil Au R1.2/1.3, 300 mesh). Then, the grids were blotted for 3.0 s or 3.5 s with 100% humidity at 4 °C, and flash-frozen in liquid ethane cooled by liquid nitrogen with Vitrobot (Mark IV, Thermo Fisher Scientific).

Micrographs were collected using a Gatan K2 Summit detector (Gatan Company) mounted on a Titan Krios electron microscope (FEI Company) operating at 300-kV and equipped with a GIF Quantum energy filter (slit width 20 eV). Micrographs were recorded in the super-resolution mode with a normal magnification of 130,000x, resulting in a calibrated pixel size of 0.53 Å. Each stack of 32 frames was exposed for 8 seconds, with an exposing time of 0.25 second per frame. The total dose rate was about 5.3 counts/sec/physical-pixel (~4.7 e-/sec/Å^2^) for each stack. AutoEMation was used for the fully automated data collection(2). All 32 frames in each stack were aligned and summed using the whole-image motion correction program MotionCor2 (3) and binned to a pixel size of 1.06 Å. The defocus value of each image was set from −1.5 to −2.0 μm and was determined by Gctf (4).

### Cryo-EM Image processing and calculation

1,338,452 particles were automatically picked by Gautomatch (developed by Kai Zhang, http://www.mrc-lmb.cam.ac.uk/kzhang/Gautomatch/) from 5,332 micrographs. Two rounds of 2D classification was performed using RELION 3.0(5, 6), resulting 923,886 particles. A small subset of those selected particles was used to generate the initial model of CSW complex with C2 symmetry by RELION 3.0. Two rounds of guided multireference classification procedure were applied to these selected particles with C2 symmetry using the program RELION 3.0 *(SI Appendix,* Fig. S2). Details of this modified procedure were previously described in the manuscript reporting the cryo-EM structure of the human spliceosomal C* complex(7). The 3D volumes of the CSW complex and five bad classes were used as initial references *(SI Appendix,* Fig. S2). These six references were low-pass filtered to 40 Å. After one round of global classification and one round local classification, 347,925 particles that belong CSW complex were separated from the bad ones. With the coordinates of these 347,925 good particles as additional input for Gautomatch, one round of exclusive particle picking was performed. With this strategy, 1,313,534 particles with different coordinates compared with the 347,925 good particles were picked. After the same procedure, these 1,313,534 particles contribute 259,603 additional particles, all together, 607,528 particles were generated for further calculation. After additional round of multi-reference classification on the same 607,528 particles with pixel size: 2.12 Å, 397,055 particles were classified out and these particles were re-centered and re-extract with pixel size 1.06 Å (*SI Appendix,* Fig. S2).

For these 397,055 particles with pixel size 1.06 Å, one round multi-reference local 3D classification with C2 symmetry was performed using two references low-pass filtered to different resolution (*SI Appendix*, Fig. S2). One class with 212,774 particles was generated. The C2 symmetry of the CSW complex is not perfect, these particles were expanded to 425,548 particles by applying a C2 operator in RELION, and further focused on one protomer of the CSW complex with C1 symmetry. With additional round of 3D classification and auto-refinement by applying soft mask, the focused protomer of the CSW complex yielded a reconstruction at an average resolution of 3.2 Å. The WDR41 and C9ORF72 region of this protomer were also further calculated to a relatively higher resolution by applying soft mask for these regions, the resolution was improved to 3.4 and 3.3 Å, respectively (*SI Appendix,* Fig. S2, *S3A-B;* Table S1).

The angular distributions of the particles used for the final reconstruction of the CSW complex are reasonable (*SI Appendix,* Fig. S3*C*), and the refinement of the atomic coordinates did not suffer from severe over-fitting (*SI Appendix,* Fig. S3*D*). The resulting density maps display clear features for the secondary structural elements and side chains of the CSW complex in the core region.

Reported resolutions were calculated on the basis of the FSC 0.143 criterion, and the FSC curves were corrected with high-resolution noise substitution methods(8). Prior to visualization, all density maps were corrected for the modulation transfer function (MTF) of the detector, and then sharpened by applying a negative B-factor that was estimated using automated procedures(9). Local resolution variations were estimated using ResMap(10).

### Model Building and refinement

The atomic coordinates of the CSW complex was generated by combining homology modelling and *de novo* model building. The structure of Longin and DENN domains of C9ORF72 and SMCR8 were generated according to the structure of DENND1b(11). For WDR41, the atomic coordinates of crystal structure of WD40 protein FBXW7 (PDB code: 5V4B(12)) was served as the initial template. These structures were docked into the density map and manually adjusted and built by by COOT(13). Sequence assignment was guided mainly by bulky residues such as Phe, Tyr, Trp and Arg. Unique patterns of sequences were exploited for validation of residue assignment.

The final overall models of the CSW complex were refined against the 3.2-Å map, using PHENIX(14) in real-space. Overfitting of the model of the CSW complex was monitored by refining the model in one of the two independent maps from the gold-standard refinement approach, and testing the refined model against the other map (15) *(SI Appendix,* Fig. S3*D*). The structures of the human CSW complex were validated through examination of the Molprobity scores and statistics of the Ramachandran plots *(SI Appendix,* Table S1). Molprobity scores were calculated as described (16).

### Pulldown assay

For the pull-down assay, indicated versions of His8-tagged SMCR8-C9ORF72 (12.5 μM) and indicated versions of untagged-WDR41(20 μM) were incubated with Ni-NTA agarose beads in incubation buffer (150mM NaCl, 25mM Tris pH 8.0, 0.05% NP40, and 5% glycerol) at 4 °C for 45 min. After rinsed with incubation buffer for three times, the resin was resuspended in 1x SDS loading buffer and boiled at 95 °C for 5 mins before applied to SDS-PAGE analysis.

### GTPase activity assay

GTPase activity assays were carried out using GTPase-Glo assay kit (Promega, V7681). Briefly, preparing 2X GTPase solution containing 1mM DTT and Rabs in buffer A (50 mM Tris-HCl, pH 7.5, 50 mM NaCl, 20 mM EDTA, and 5 mM MgCl_2_). The concentration of Rab in this solution is 2-fold of the final concentration. Then, diluting GAP to desired concentration in buffer A containing 10 μM GTP, and dispense 5 μl into each well of a 384-well plate (Corning, 3570). To initiate the GTPase reaction, 5 μl 2X GTPase solution was added into each well. The total GTPase reaction volume is 10 μl. The plate was incubated at 25 °C for 120 min. Then, 10 μl of reconstituted GTPase-Glo reagent was added to the wells to terminate the GTP hydrolysis reactions and the plate was incubated at 25°C for 30 min with shaking. Then, 20 μl of the Detection Reagent (from the kit) was added to all the wells. The luminescence signal was measured using BioTek Synergy2. Noteworthily, the higher the luminescence is, the less the GTP is consumed by GTPases.

### Nucleotide exchange assay

Purified Rabs were loaded with MANT-GDP (Thermo Fisher, M12414) in the presence of 10 mM EDTA pH 8.0 and 2-fold molar excess of MANT–GDP at 4 °C overnight. Loading reaction was terminated by adding MgCl_2_ to 50 mM and the Rab8a-MANT-GDP complex was further purified using Superdex75 10/300 (GE Healthcare) in 25 mM Tris-HCl pH 8.0, 150 mM NaCl, 5 mM MgCl_2_ and 2 mM DTT. For the nucleotide exchange assay, 1.6 μM Rab GTPase-MANT-GDP complex were pre-incubated with 0 and 300 nM of indicated C9ORF72-SMCR8 subcomplex for 30 minutes. After baseline stabilization, the nucleotide exchange reaction was triggered by the addition of GDP to a final concentration of 0.1 mM. Releasing of MANT-GDP was recorded by monitoring the decrease in fluorescence emission at 448 nm (excited at 360 nm) in intervals of 15 s at 25 °C for 3600-second. Data were collected using 384-well plate (Corning, 3701) in BioTek Synergy2.

**Fig. S1.**
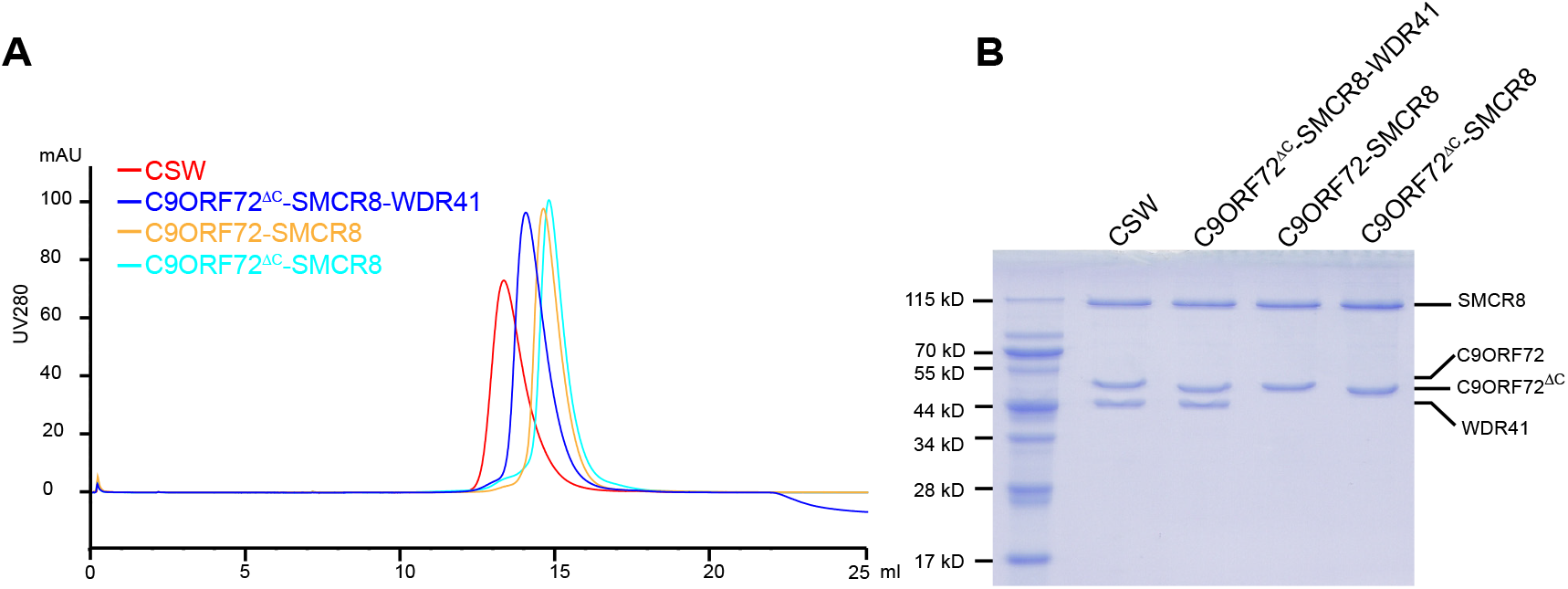
Gel filtration profiles of different complexes. **A.** The elution profiles of different complexes in Superose 6 10/300 GL column are shown here. Red: CSW, blue: C9ORF72^ΔC^-SMCR8-WDR41, orange: C9ORF72-SMCR8, cyan: C9ORF72^ΔC^-SMCR8. C9ORF72^ΔC^-SMCR8-WDR41 was eluted earlier than C9ORF72-SMCR8 due to C9ORF72^ΔC^-SMCR8-WDR41 is more elongated than C9ORF72-SMCR8. **B.** The peak fractions from gel filtrations are visualized with the Coomassie blue-stained SDS-PAGE.

**Fig. S2.**
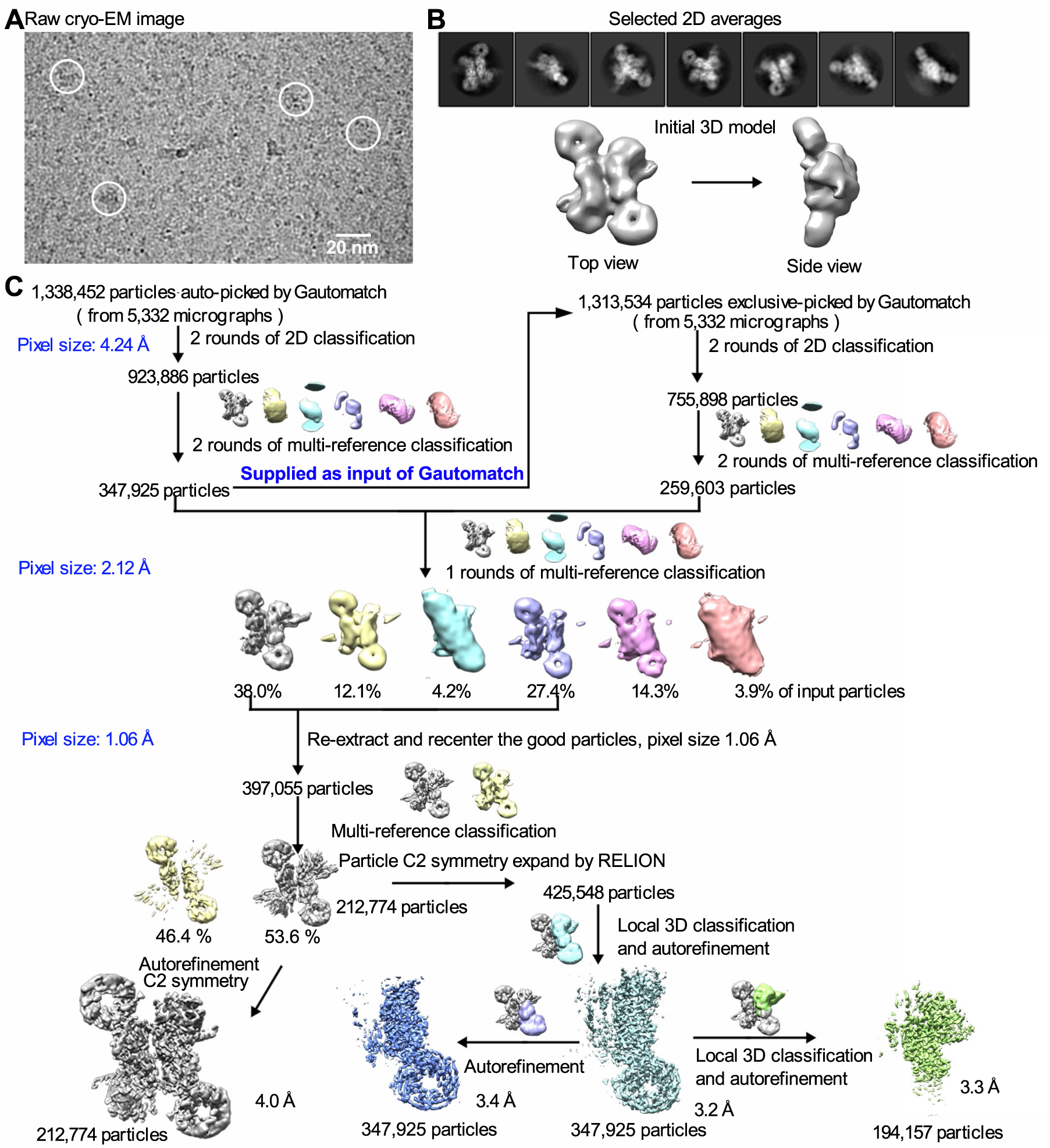
A flow chart for the cryo-EM data processing and structure determination of the human C9ORF72-SMCR8-WDR41 complex. **A.** A representative cryo-EM micrograph of the final CSW complex. Scale bar, 20 nm. **B.** Selected 2D classification averages and an initial model. **C.** Based on the FSC value of 0.143, the final reconstruction has an average resolution of 3.2 Å for one protomer of the CSW complex. Please refer to the Method for the detailed description.

**Fig. S3.**
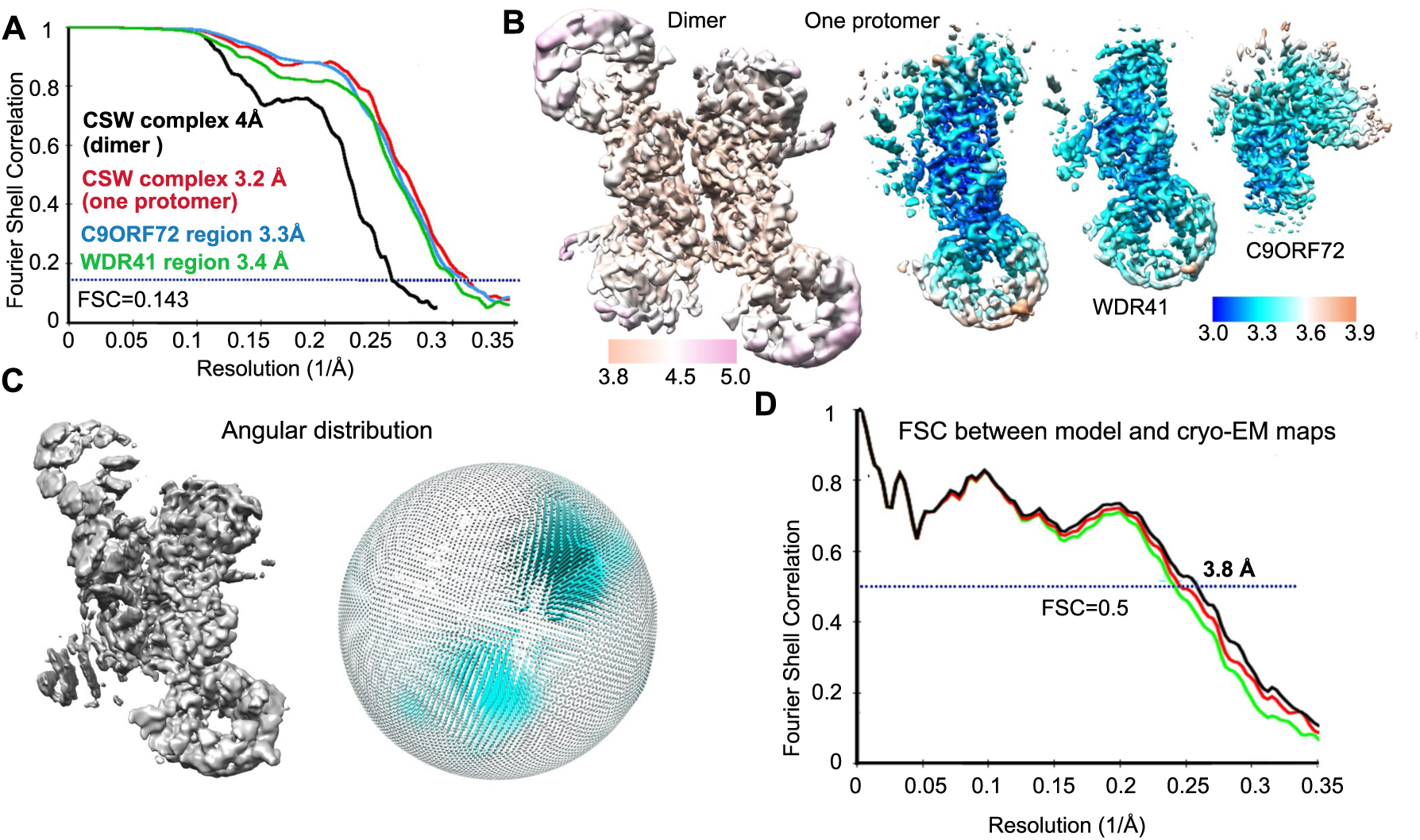
Cryo-EM analysis of the human C9ORF72-SMCR8-WDR41 complex. **A.** The average resolutions of the map for CSW complex are estimated to be 3.2 Å for one protomer based on the FSC criterion of 0.143. The average resolutions for WDR41 region and C9ORF72 region are estimated to be 3.4 and 3.3 Å, respectively. B. The local resolutions are color-coded for different regions of the CSW complex, including two sub-regions of WDR41 and C9ORF72. **C.** Angular distribution of the particles used for the reconstruction of the human CSW complex. Each cylinder represents one view, and the height of the cylinder is proportional to the number of particles for that view. **D.** The FSC curves of the final refined model of the CSW protomer versus the overall map that it was refined against (black); of the model refined in the first of the two independent maps used for the gold-standard FSC versus that same map (red); and of the model refined in the first of the two independent maps versus the second independent map (green). The generally similar appearances between the red and green curves indicate that the refinement of the atomic coordinates did not suffer from severe over-fitting. 3.8 Å indicates the resolution of the black curve at FSC=0.5.

**Fig. S4.**
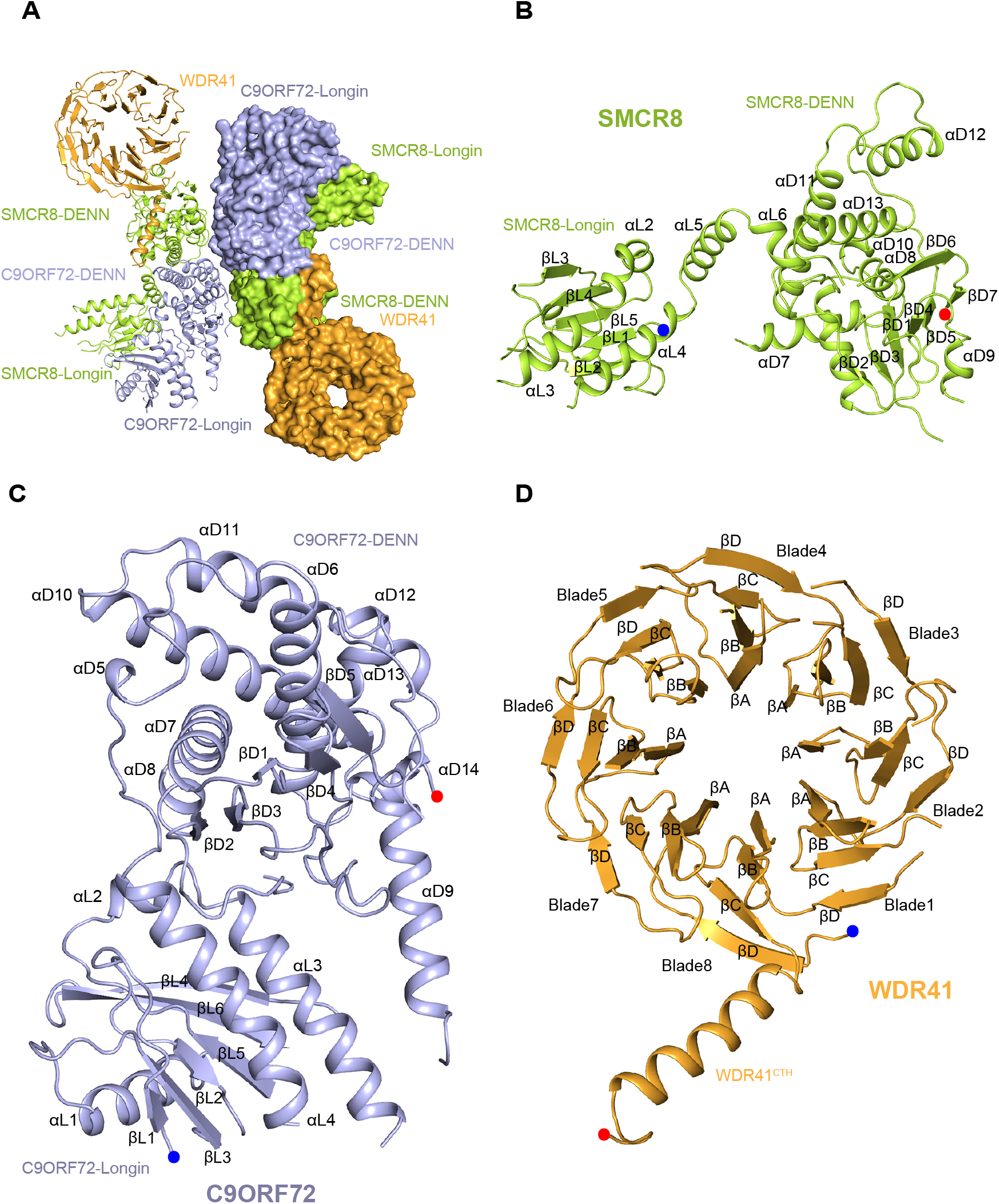
The architecture of each subunit of CSW. **A.** The overall structure of CSW. **B.** Structure of SMCR8. **C.** Structure of C9ORF72. **d.** Structure of WDR41. The secondary structures are labeled. The blue and red dots indicated the N-termini and C-termini of proteins.

**Fig. S5.**
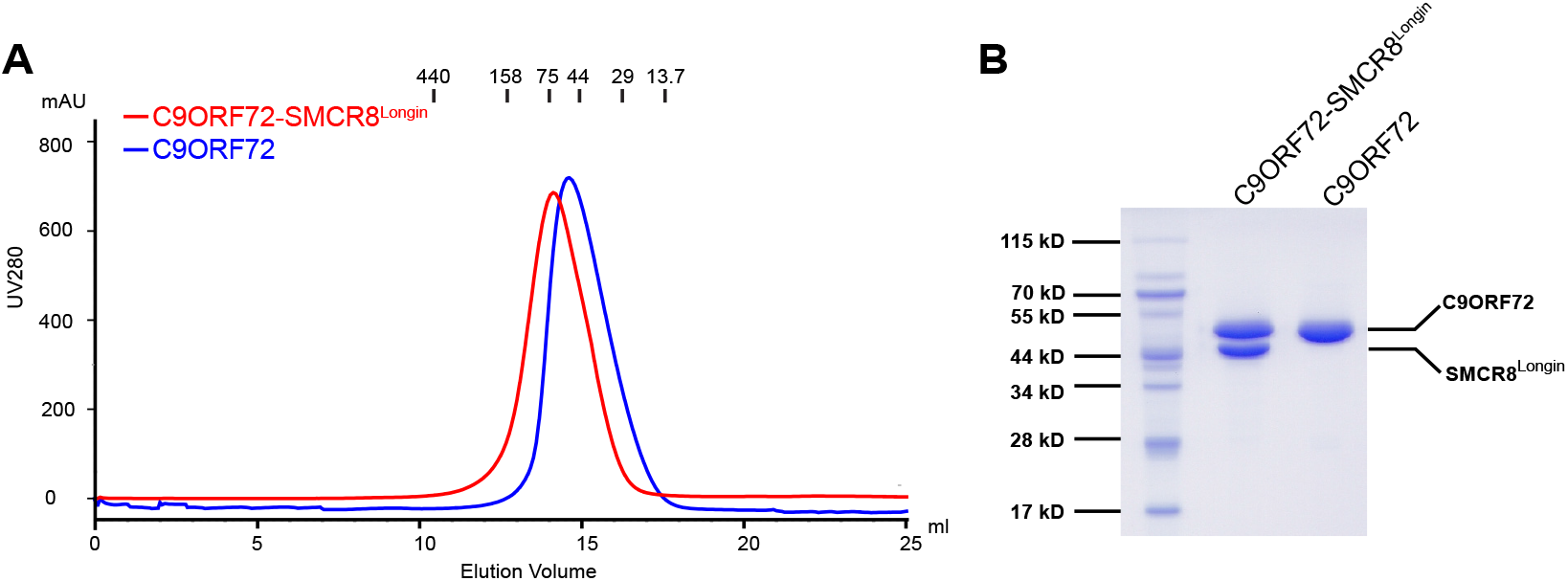
Gel filtration profiles of C9ORF72 and C9ORF72-SMCR^Longin^. **A.** The elution profiles of different complexes in the Superdex 200 10/300 GL column are shown here. Red: C9ORF72-SMCR^Longin^; blue: C9ORF72. The molecular weight standers are indicated on top. **B.** The peak fractions from panel (A) were visualized with the Coomassie blue-stained SDS-PAGE.

**Fig. S6.**
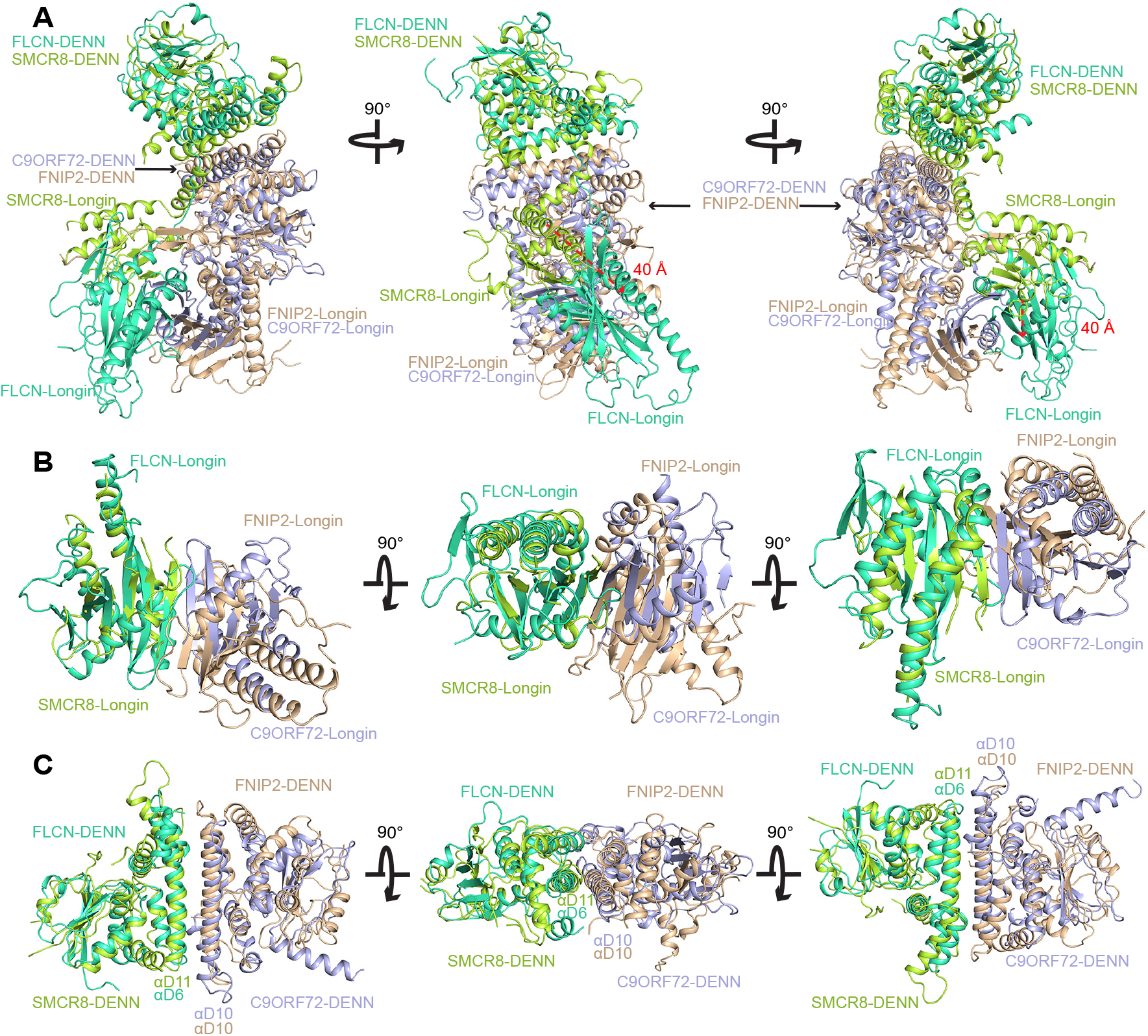
The architecture of C9ORF72-SMCR8 complex protomer. **A.** Different views of the superimposition of the C9ORF72-SMCR8 complex protomer and the FLCN-FNIP2 complex. In the middle and right panels, the red arrow indicates the distance between SMCR8^Longin^ and FLCN^Longin^ when the DENN dimers of two complexes are overlapped. (**B, C**) Comparison of Longin dimers (**B**) and DENN dimers (**C**) of C9ORF72-SMCR8 complex with those of FLCN-FNIP2 dimer. Three views are presented here. The helixes mediated the formation of DENN dimer are labeled in panel (**C**).

**Fig. S7.**
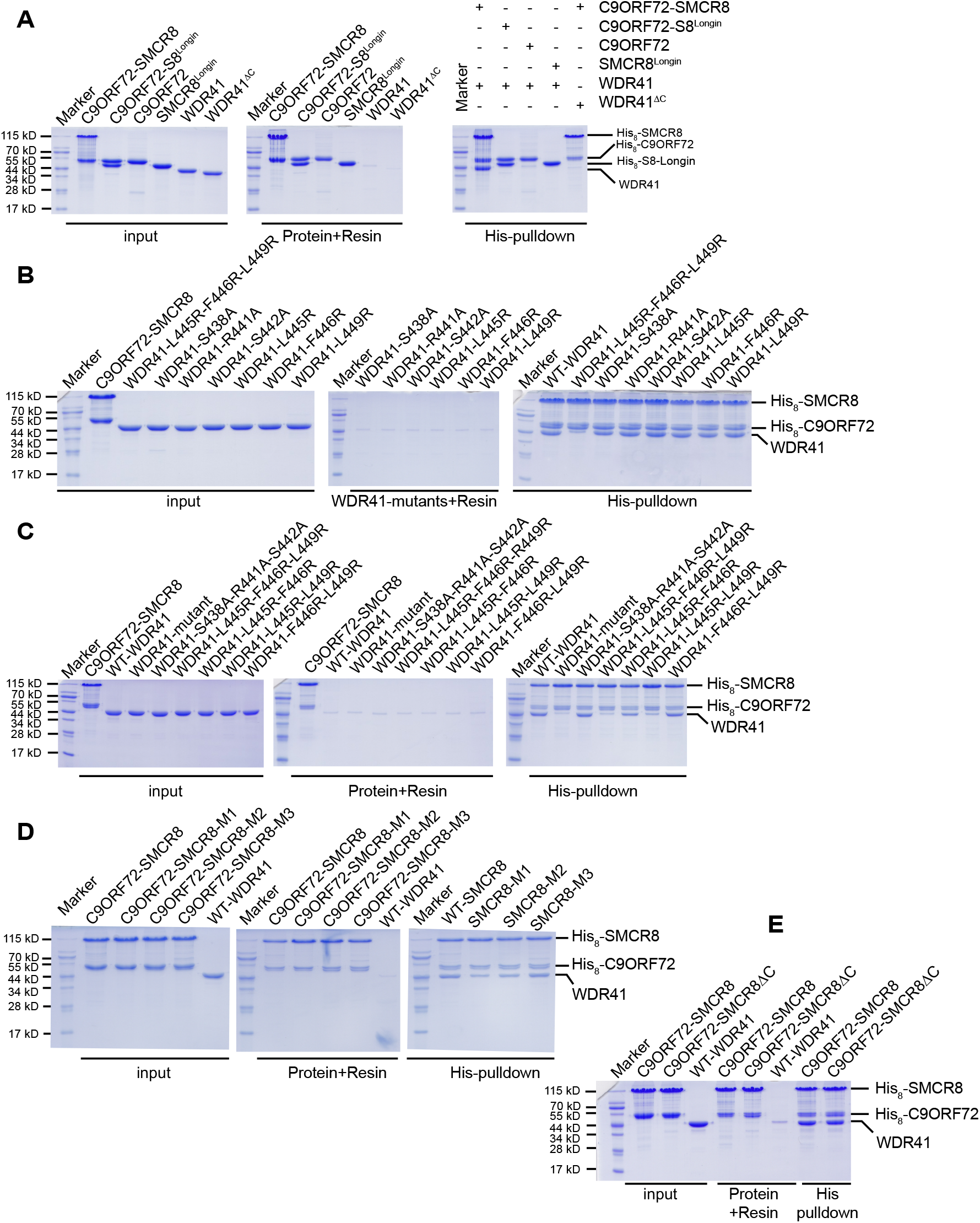
Contributions of key residues to the interface of SMCR8 and WDR41. **A.** Full profile of pulldown assay of Fig. 4B. The labels are same to Figure 4B. Protein markers are labeled at left (unit: kD). **B.** Effect of single mutations of WDR41 on the bind to SMCR8 was assessed by pull down assay. The mutated residues are labeled. **C.** Full profile of pulldown assay of Fig. 4C. The labels are same to Figure 4C. **D.** Full profile of pulldown assay of Fig. 4D. The labels are same to Figure 4D. **E.** Effect of βD6 of SMCR8 on the binding to WDR41was assessed by pulldown assay. The labels are same to Figure 4. C9ORF72-SMCR8^ΔC^: C9ORF72-SMCR8 (SMCR8 with 923-937 deleted). WT-WDR41: wild-type WDR41.

**Fig. S8.**
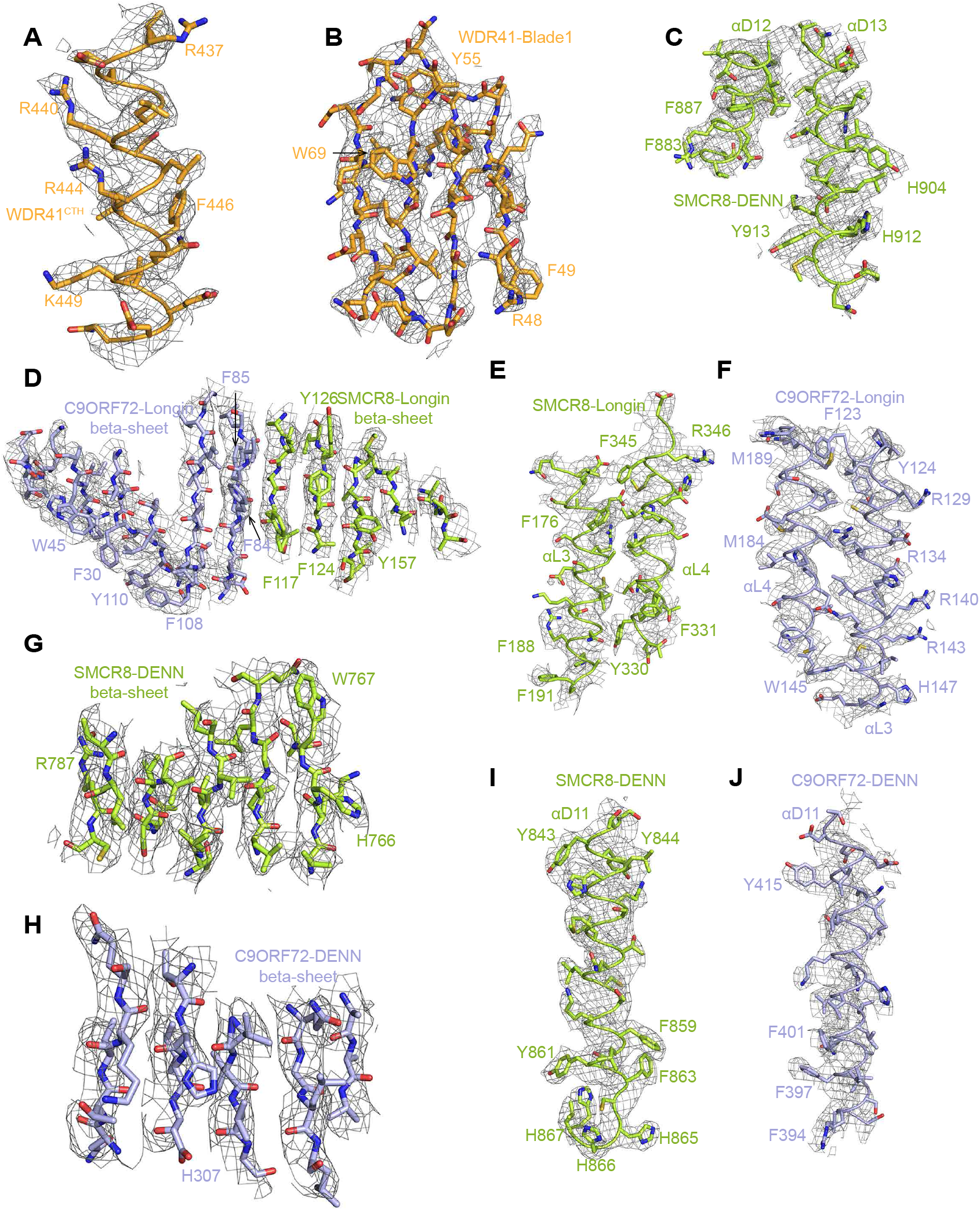
EM map for the indicated region. The core regions of the model fit well with the EM density. (**A-J**) Different regions of the CSW protomer are shown as indicated. The bulky residues are labeled.

**Fig. S9.**
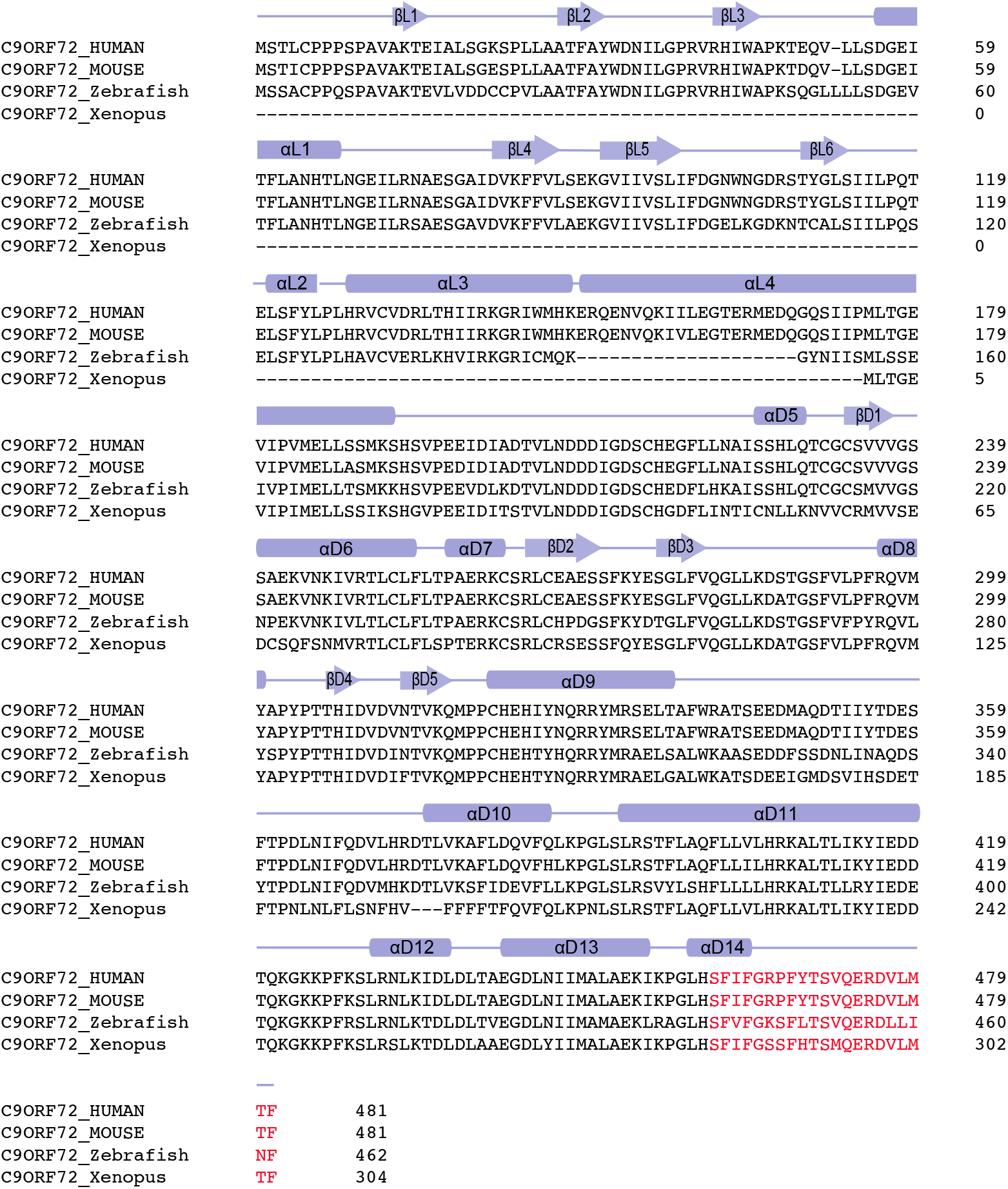
Sequence alignment of C9ORF72. HUMAN: *Homo sapiens;* Mouse: *Mus musculus;* Zebrafish: *Danio rerio;* Xenopus: *Xenopus tropicalis;* The deleted residues are indicated as red. The secondary structures are labeled on top of the sequences.

**Fig. S10.**
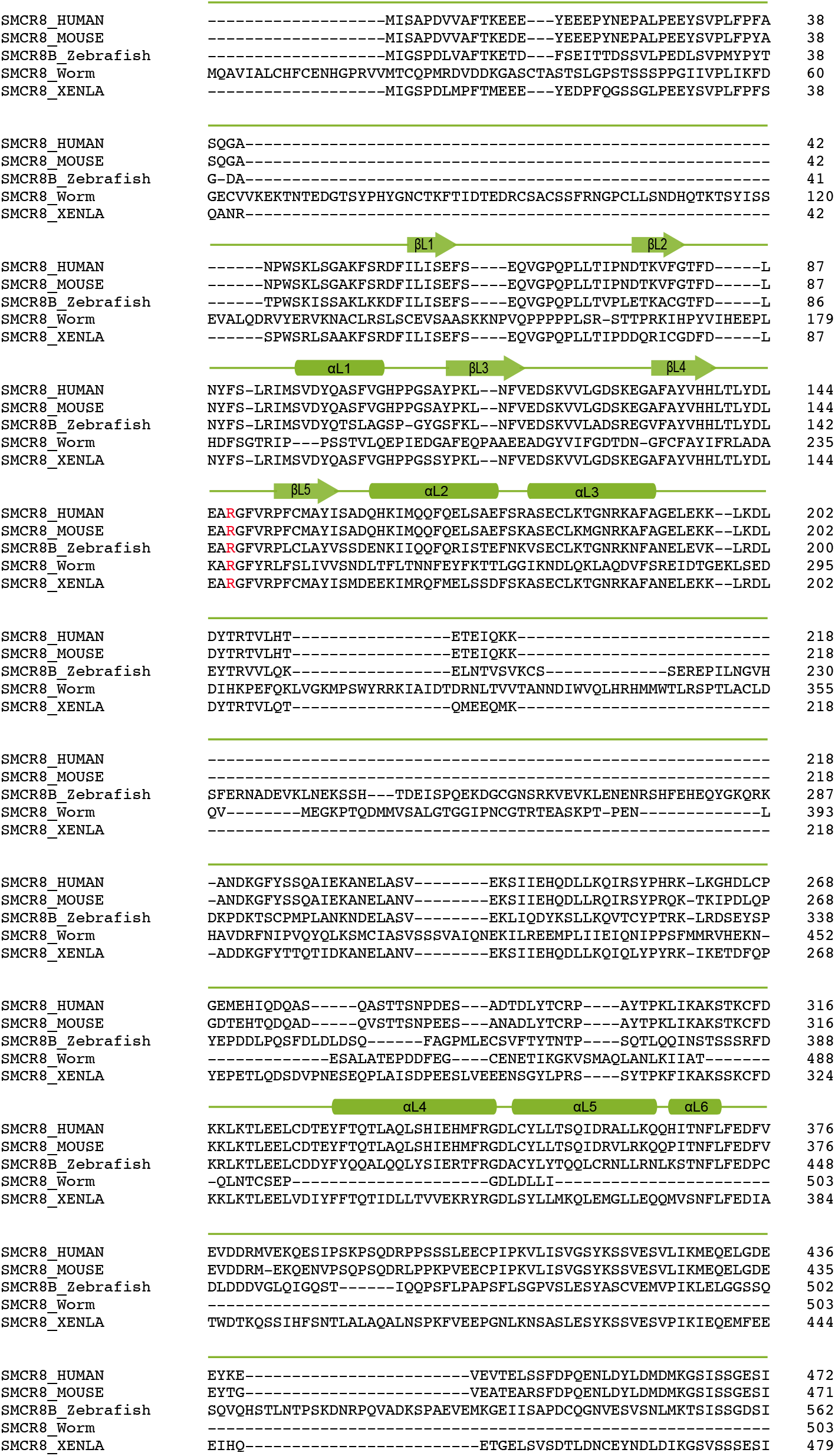
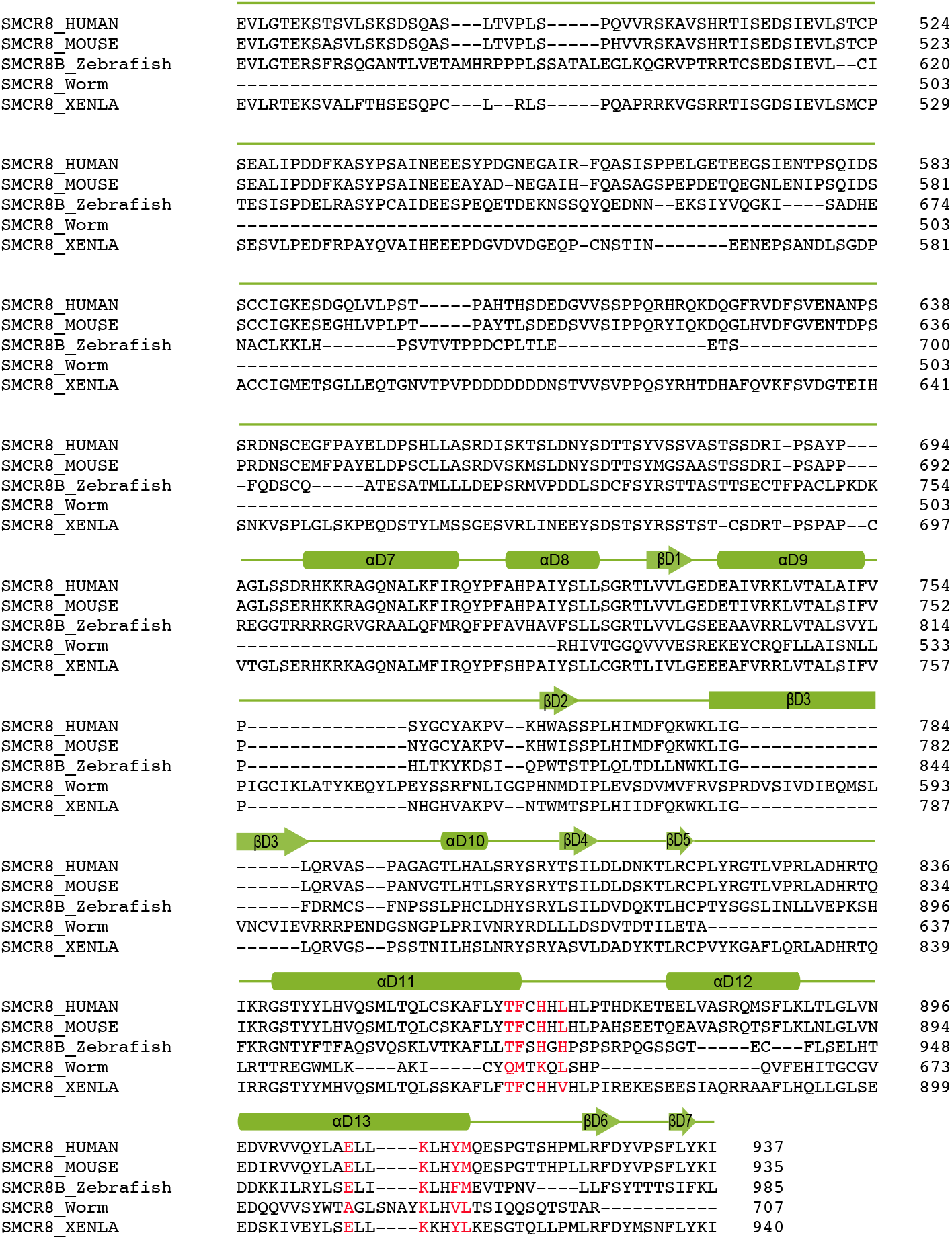
Sequence alignment of SMCR8. HUMAN: *Homo sapiens;* Mouse: *Mus musculus;* Zebrafish: *Danio rerio;* Worm: *Caenorhabditis elegans;* Xenopus: *Xenopus tropicalis;* The mutated residues are indicated as red. The secondary structures are labeled on top of the sequences.

**Fig. S11.**
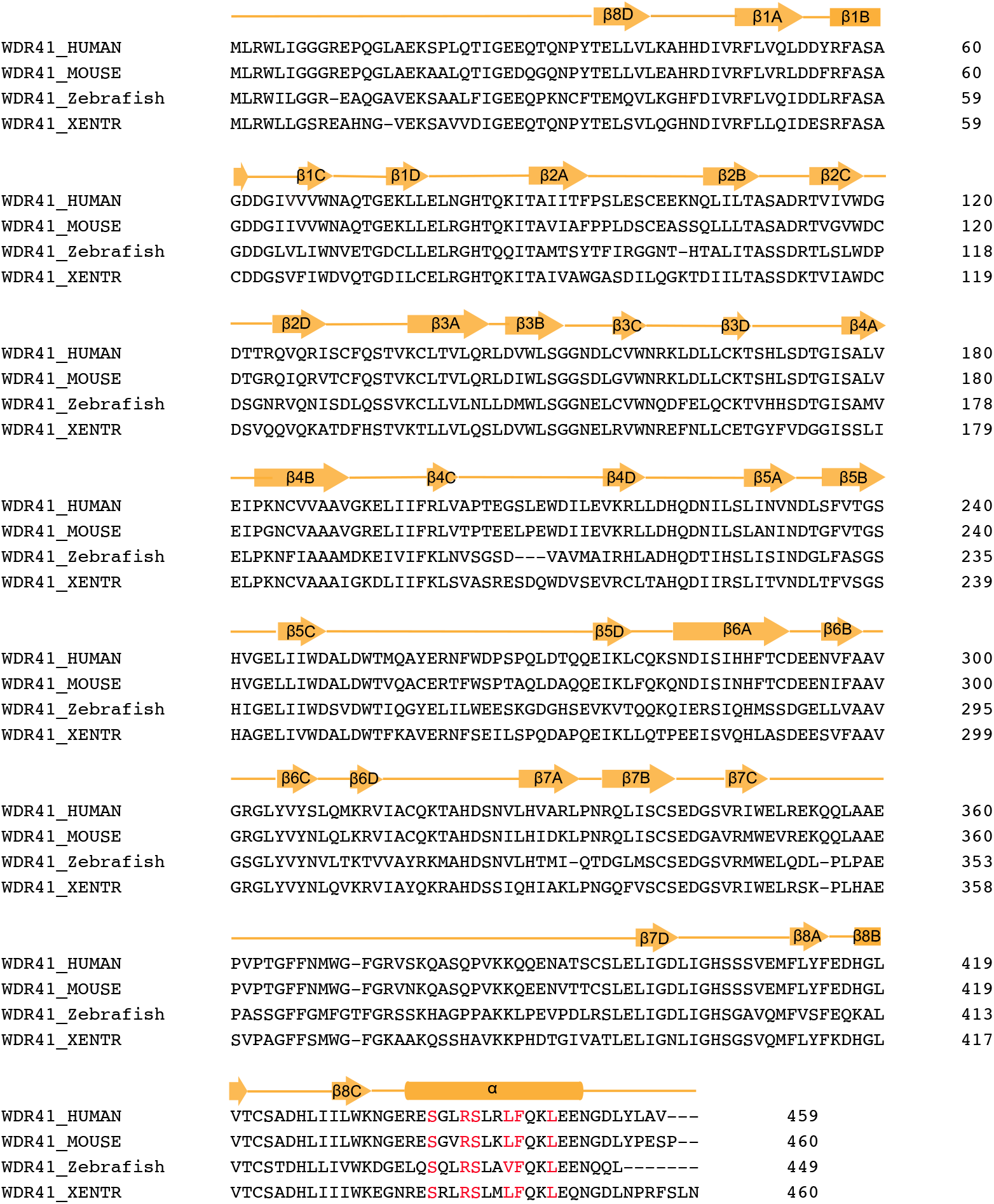
Sequence alignment of WDR41. HUMAN: *Homo sapiens;* Mouse: *Mus musculus;* Zebrafish: *Danio rerio;* Xenopus: *Xenopus tropicalis;* The mutated residues are indicated as red. The secondary structures are labeled on top of the sequences.

**Table S1.**
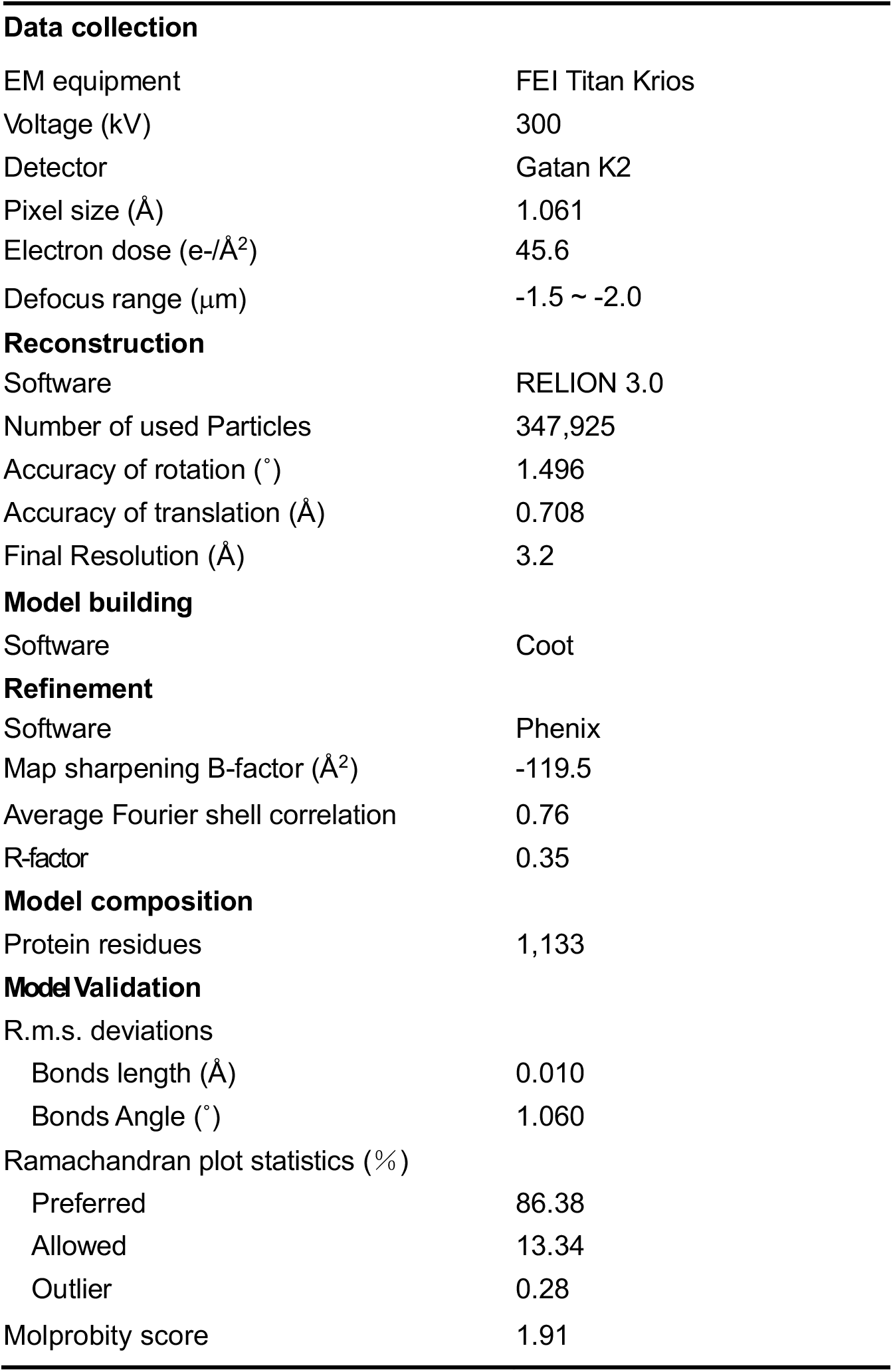
Cryo-EM data collection and refinement statistics.

